# Endothelial RIPK3 minimizes organotypic inflammation and vascular permeability in ischemia-reperfusion injury

**DOI:** 10.1101/2024.11.25.625188

**Authors:** Charmain F. Johnson, Christopher M. Schafer, Kathryn Y. Burge, Brian G. Coon, Hala Chaaban, Courtney T. Griffin

## Abstract

Recent studies have revealed a link between endothelial receptor-interacting protein kinase 3 (RIPK3) and vascular integrity. During mouse embryonic development, hypoxia can trigger elevated endothelial RIPK3 that contributes to lethal vascular rupture. However, it is unknown whether RIPK3 regulate endothelial barrier function in adult vasculature under hypoxic injury conditions such as ischemia-reperfusion (I/R) injury. Here we performed inducible genetic deletion of endothelial *Ripk3* (*Ripk^iECKO^*) in mice, which led to elevated vascular permeability in the small intestine and multiple distal organs after intestinal I/R injury. Mechanistically, this vascular permeability correlated with increased endothelial secretion of IL-6 and organ-specific expression of VCAM-1 and ICAM-1 adhesion molecules. Circulating monocyte depletion with clodronate liposomes reduced permeability in organs with elevated adhesion molecules, highlighting the contribution of monocyte adhesion and extravasation to *Ripk^iECKO^* barrier dysfunction. These results elucidate mechanisms by which RIPK3 regulates endothelial inflammation to minimize vascular permeability in I/R injury.

**GRAPHICAL ABSTRACT:** 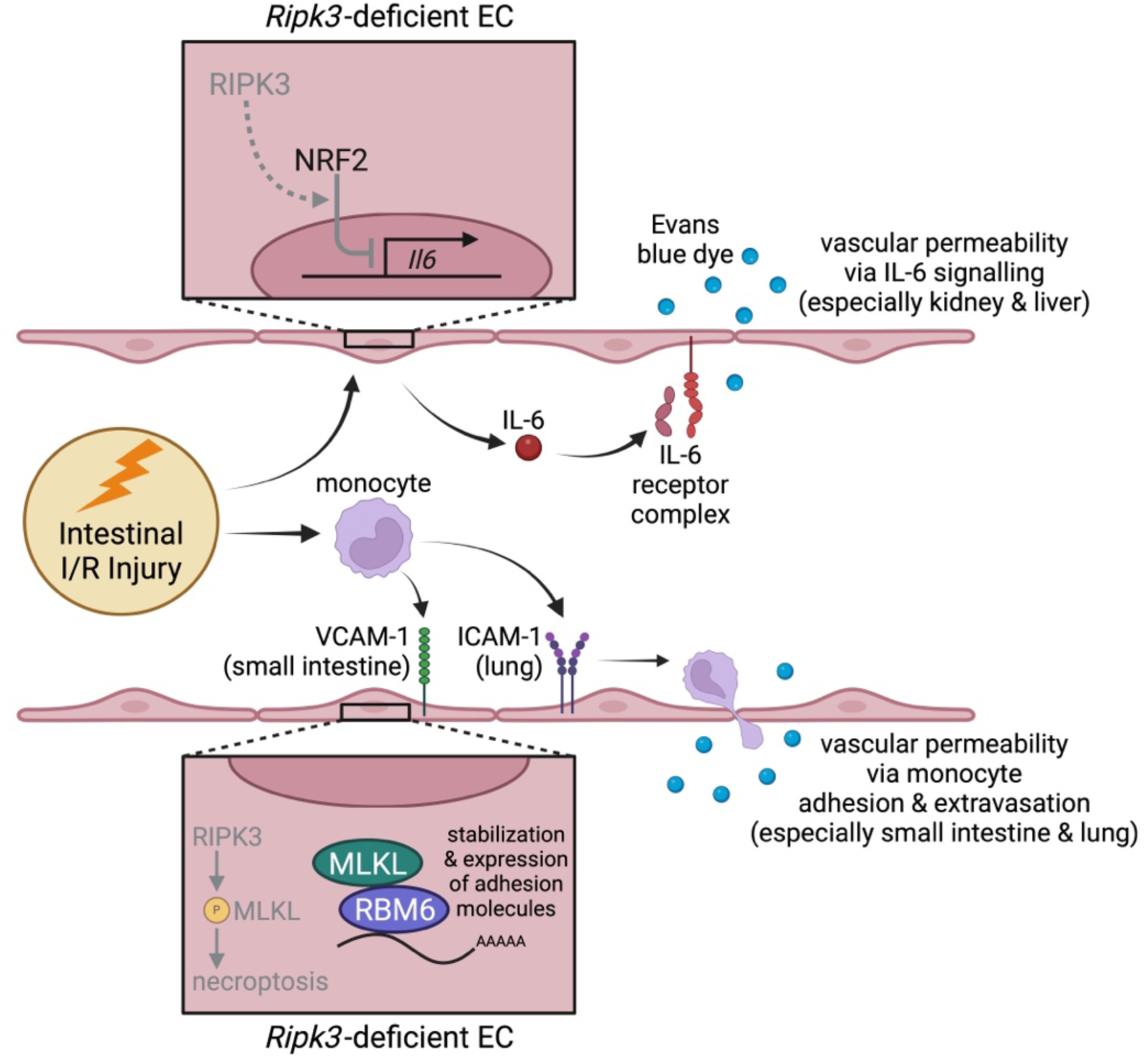

## INTRODUCTION

Receptor-interacting protein kinase 3 (RIPK3) is a cytoplasmic serine-threonine protein kinase that was initially identified as a key component of the necroptosis cell death pathway^1–3^ and has a growing list of non-necroptotic functions that contribute to inflammatory diseases^4^. Our lab has identified several non-necroptotic roles for RIPK3 in endothelial cells (ECs). For example, we found that elevation of endothelial RIPK3 in midgestation mouse embryos promotes lethal vascular rupture without activating its critical downstream necroptosis component, mixed lineage kinase domain-like (MLKL)^5^. This finding is consistent with studies in neonatal and juvenile mice, in which RIPK3 destabilizes EC adherens junctions to impair developmental and pathological ocular angiogenesis^6^. It has also been reported that RIPK3 utilizes both necroptotic and non-necroptotic mechanisms to promote vascular permeability that facilitates metastatic tumor cell extravasation out of the blood stream in adult mice^7,8^. These studies collectively indicate that excessive endothelial RIPK3 promotes detrimental vascular permeability through multiple mechanisms.

Endothelial RIPK3 protein expression is naturally high in midgestation mouse embryos at the stage in which they are most hypoxic, prior to the establishment of the fetal-placental circulation^9,10^. We reported that hypoxia can trigger *Ripk3* transcription in embryonic ECs, particularly in conjunction with suppression of the inhibitory chromatin remodeling enzyme chromodomain helicase DNA binding protein 4 (CHD4)^5^. Hypoxia has also been shown to contribute to elevated RIPK3 expression in cultured human hepatocyte and murine neuronal cell lines^11,12^. Moreover, several mouse models of ischemic diseases are ameliorated on a *Ripk3^-/-^* background^13,14,15^, suggesting that hypoxia-induced RIPK3 expression drives tissue pathology. However, whether hypoxia triggers RIPK3 upregulation in adult ECs and triggers RIPK3-mediated vascular permeability has not yet been explored.

Intestinal ischemia-reperfusion (I/R) injury is a clinically important patho-physiological process resulting from a broad range of surgical and non-surgical causes, including acute mesenteric ischemia^16,17^, shock^18^, neonatal necrotizing enterocolitis^19^, trauma^20^, and intestinal transplantation^21^. During the ischemic phase of intestinal I/R injury, blood flow to the intestines is reduced due to obstruction of mesenteric arteries supplying blood to the small and large intestines^17,22^. The current therapeutic approach for addressing intestinal ischemia is to resolve the obstruction and restore blood flow to the intestinal tissue as soon as possible^23^. Although reperfusion of the tissue is crucial, this process can trigger further pathological damage to the intestine through generation of reactive oxygen species that can mediate cell death and breakdown of the local mucosal and vascular barriers. Bacterial leakage and circulating factors (e.g., inflammatory cytokines) produced in the reperfused intestinal region can then be transported in the bloodstream and cause damage to organs such as the heart, liver, and lungs that are remote/distal to the original injury site^23–26^. Because intestinal I/R injury triggers transient tissue hypoxia and both local and distal vascular permeability, we identified this as an ideal model for investigating the hypoxic upregulation of RIPK3 in ECs and its subsequent effects on vascular integrity in adult mice.

In this study, we investigated the role of endothelial-specific RIPK3 in intestinal I/R injury using genetic, surgical, and molecular approaches. We found that hypoxia and I/R injury upregulate RIPK3 expression in adult ECs. Interestingly, this upregulation is protective to the vasculature, since deletion of endothelial *Ripk3* promotes enhanced inflammation and vascular permeability both in the small intestine and in some remote organs after I/R injury. Therefore, endothelial RIPK3 plays a beneficial role in the context of intestinal I/R injury by contributing to organotypic vascular integrity.

## RESULTS

### Deletion of endothelial Ripk3 in established blood vessels causes vascular hyperpermeability following intestinal I/R injury

Because we previously reported that embryonic endothelial RIPK3 expression can be elevated by hypoxia^5^, we questioned whether endothelial RIPK3 also increases after ischemic challenge in adult mice. We addressed this by assessing RIPK3 expression in *en face* mesenteric arteries following intestinal ischemia and 24 hr reperfusion. Because commercial RIPK3 antibodies give nonspecific immunostaining in our hands, we utilized a RIPK3-GFP reporter mouse^27^ and immunostained for GFP. We found significantly elevated RIPK3/GFP expression in mesenteric ECs after one hour of ischemic clamping, and this elevated expression was maintained even after 24 subsequent hours of reperfusion (**Supplemental Figure S1A, B**). We next examined *Ripk3* mRNA expression in the MS1 adult murine EC line cultured in glucose-free media under hypoxic (1% O_2_) versus normoxic conditions and found a significant increase in *Ripk3* expression at 12 hr after oxygen deprivation (**Supplemental Figure S1C**), suggesting that hypoxia drives RIPK3 expression in adult murine ECs.

Since elevated endothelial RIPK3 is associated with blood vessel permeability and rupture^5,6,8^, we next explored the role of endothelial RIPK3 in I/R injury-induced vascular permeability. We generated *Ripk3^fl/fl^;Cdh5(PAC)-Cre^ERT2^*mice (hereafter called *Ripk3^iECKO^*) to achieve tamoxifen-inducible *Ripk3* knockout in ECs, as we previously described^28,29^. Tamoxifen was administered to 8-week-old littermate control (*Ripk3^fl/fl^*; hereafter called *Ripk3^WT^*) and *Ripk3^iECKO^* mice, and four weeks later we detected no *Ripk3* mRNA expression in ECs isolated from *Ripk3^iECko^*lungs (**Supplemental Figure S2A, B**). Notably, we found no difference in *Ripk3* expression in leukocytes collected from the peripheral blood of *Ripk3^WT^* and *Ripk3^iECKO^* mice under the same induction scheme (**Supplemental Figure S2C**). We also used a *ROSA^mT/mG^*Cre reporter mouse^30^ to assess the endothelial specificity of the *Cdh5(PAC)-Cre^ERT2^* transgene and found that Cre activity (GFP expression) was restricted to ECs and was not detected in tissue resident leukocytes (**Supplemental Figure S2D**) or circulating blood cells (**Supplemental Figure S2E**).

We next measured baseline vascular permeability in 12-week-old littermate *Ripk3^WT^* and *Ripk3^iECKO^* mice using an Evans blue dye (EBD) perfusion and leakage assay (**Figure 1A**). We found that vascular permeability was unchanged in small intestines, livers, and hearts between *Ripk3^WT^* and *Ripk3^iECKO^* mice under baseline conditions. However, lung and kidney vascular permeability was significantly decreased in *Ripk3^iECKO^* mice (**Figure 1B**). When EBD leakage was measured following 24 hr of I/R injury, we found that vascular permeability was elevated in the intestinal region downstream of clamped mesenteric vessels (i.e., the I/R region) in both *Ripk3^WT^* and *Ripk3^iECKO^* mice (**Figure 1C**). Surprisingly, vascular permeability was also elevated in adjacent, unclamped control and distal control intestinal regions of *Ripk3^iECKO^* mice but not in littermate *Ripk3^WT^* mice (**Figure 1C**). Additionally, we found that liver, lung, and kidney vascular permeability was elevated in *Ripk3^iECKO^* mice compared to *Ripk3^WT^* mice after I/R injury (**Figure 1D**). We found no changes in vascular permeability in sham operated mice (**Supplemental Figure S3**). In addition to quantifying EBD leakage, we also perfused mice with 70 kDa FITC-dextran after I/R injury and found that in *Ripk3^iECKO^* mice, FITC-dextran leakage was mainly visualized in the submucosal regions of the small intestine where post-capillary venules are located (**Figure 1E**, white arrows). In the liver, lung, and kidneys of *Ripk3^iECKO^* mice, FITC-dextran was mostly detected surrounding capillary networks rather than larger blood vessels (**Supplemental Figure S4**).

**FIGURE 1.**
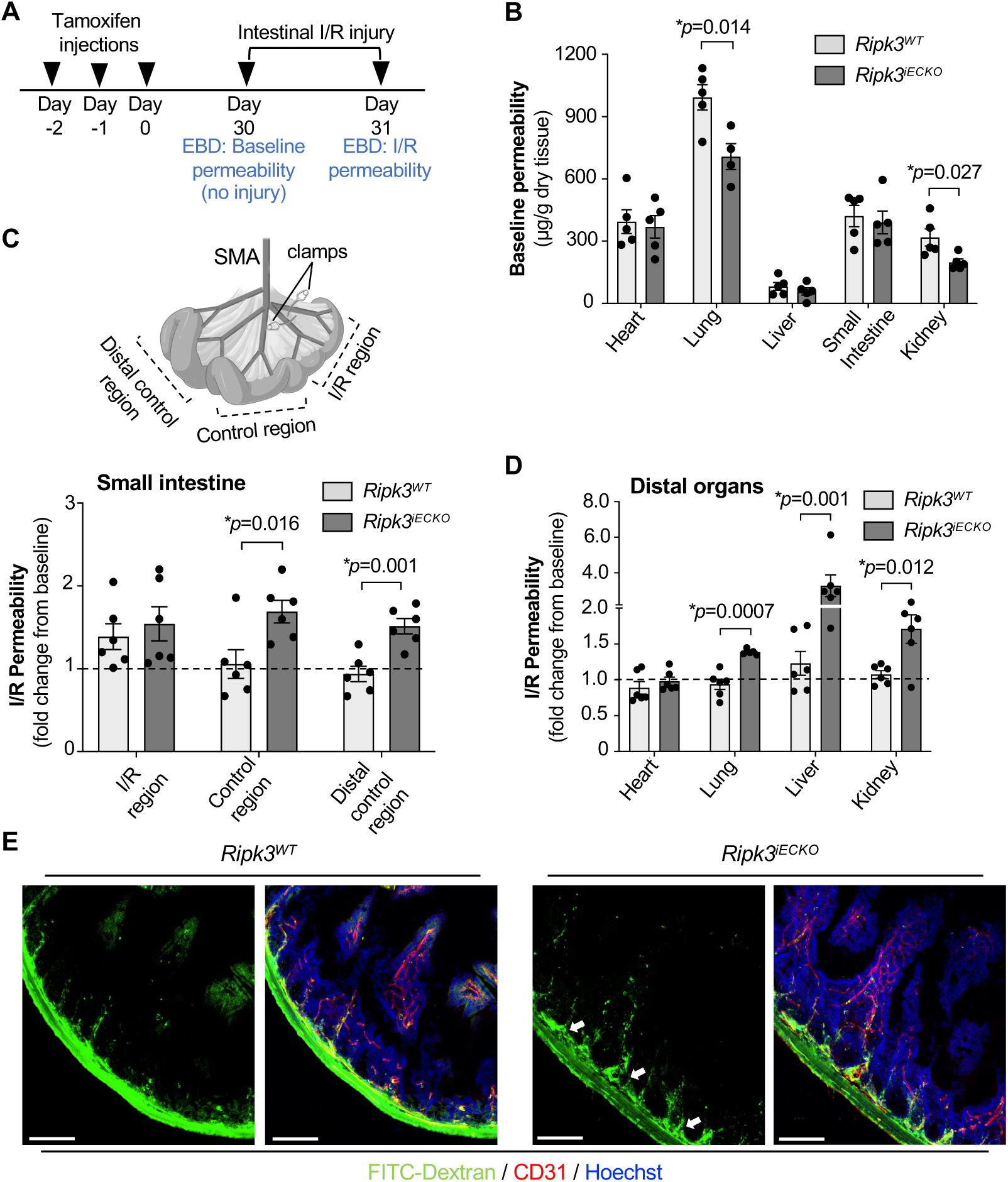
Endothelial cell (EC)-specific deletion of Ripk3 in established blood vessels causes vascular hyperpermeability following intestinal ischemia-reperfusion (I/R) injury. (**A**) Timeline for conditional deletion of EC *Ripk3* (*Ripk3^iECKO^)* in mice at 8 weeks of age and baseline or intestinal I/R injury-induced vascular permeability measurements using Evans blue dye (EBD) at 12 weeks of age. (**B**) Baseline vascular permeability measurements using EBD in *Ripk3^WT^* and *Ripk3^iECKO^* mice in the indicated organs (n=5/group). (**C**) Small intestinal vascular permeability after 24 hr intestinal I/R injury in the injury region (I/R) and control regions (control, distal control) (n=6/group); assessed regions are depicted in the schematic. (**D**) Vascular permeability in the indicated organs after 24 hr intestinal I/R injury (n=6/group). (**E**) Representative confocal Z-stack images of FITC-dextran (70kDa) permeability (white arrows) in control regions of small intestinal tissue after 24 hr intestinal I/R injury. Scale bar: 100 µm. *P* values were determined by unpaired t-tests for each organ between the two groups (**B**, **C** and **D**), and data that failed an equal variance F-test were log-transformed before analysis (**D**). \**P* <0.05 *vs*. Control. Summary data are the mean ± SE. **SMA** = superior mesenteric artery

### Loss of Ripk3 causes increased secretion of the inflammatory cytokine IL-6 from ECs

Because some cytokines secreted by activated immune cells and ECs after reperfusion injury can promote vascular permeability^23,31,32^, we analyzed several such cytokines in the serum and small intestinal tissue of *Ripk3^WT^* and *Ripk3^iECKO^* mice at baseline and after I/R injury using a multiplex immunoassay. We chose a time point of 4 hr after I/R injury (instead of 24 hr) for our analyses because many cytokine levels rise quickly after I/R injury^23^. We found that IL-6 levels were significantly elevated in the small intestine and serum of *Ripk3^iECKO^* mice after I/R injury (**Figure 2A, B**), unlike the other cytokines analyzed (**Supplemental Figure S5A-D**).

**FIGURE 2.**
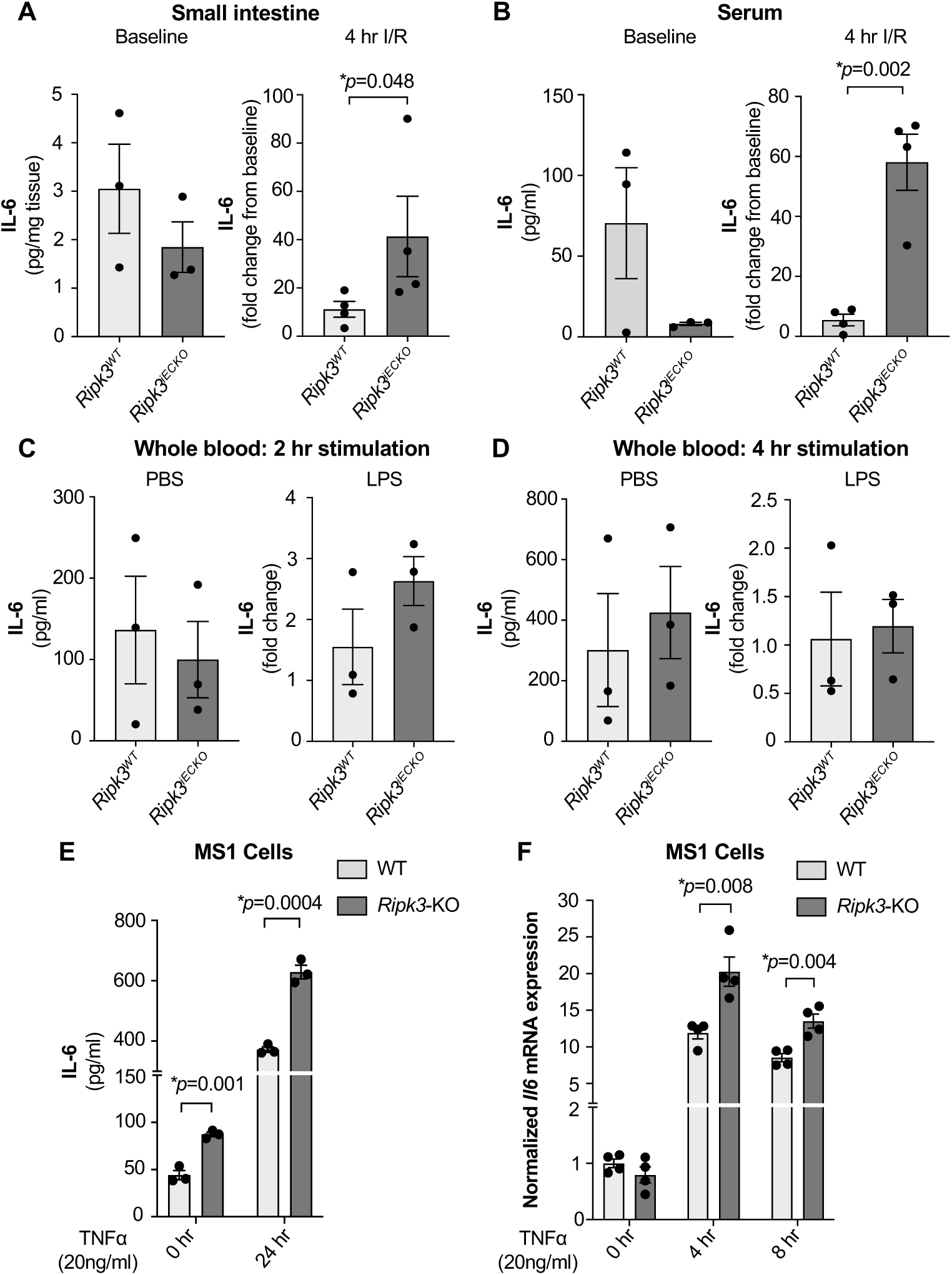
Endothelial cell (EC) secretion of the inflammatory cytokine IL-6 is elevated in the absence of Ripk3. (**A**) Baseline small intestine tissue IL-6 levels (left) and fold change in small intestine tissue IL-6 concentration following 4 hr intestinal I/R injury (right) in *Ripk3^WT^* and *Ripk3^iECKO^*mice (n=3-4/group). (**B**) Baseline serum IL-6 levels (left) and fold change in serum IL-6 concentration following 4 hr intestinal I/R injury (right) in *Ripk3^WT^* and *Ripk3^iECKO^*mice (n=3-4/group). (**C** and **D**) Plasma IL-6 levels at 2 hr (**C**) or 4 hr (**D**) after stimulating whole blood collected from *Ripk3^WT^*and *Ripk3^iECKO^* mice with phosphate-buffered saline (PBS) (left) and fold changes in IL-6 levels (from PBS stimulation) after stimulating with lipopolysaccharide (LPS) (right) (n=3/group). (**E**) IL-6 levels in cell supernatants after stimulating WT and *Ripk3*-KO MS1 ECs with 20ng/mL TNFα for 24 hr. (**F**) *Il6* transcripts in WT and *Ripk3*-KO MS1 ECs following stimulation with 20ng/mL TNFα for 0 hr, 4 hr and 8 hr. *P* values were determined by unpaired t-tests for each treatment condition between the two groups (**A**-**F**), and data that failed an equal variance F-test were log-transformed before analysis (**A** and **B**). \**P* <0.05 *vs*. Control. Summary data are the mean ± SE.

To determine the cell source of the increased IL-6 that we detected in *Ripk3^iECKO^* mice after I/R injury, we collected whole blood from *Ripk3^WT^* and *Ripk3^iECKO^* mice via cardiac punctures, stimulated blood immune cells with bacterial LPS (or PBS) for 2 or 4 hr, and measured plasma IL-6 levels. We did not observe differences in IL-6 levels secreted from stimulated blood from *Ripk3^WT^* versus *Ripk3^iECKO^* mice at either time point (**Figure 2C, D**), indicating that circulating immune cells were not the source of the elevated IL-6 in *Ripk3^iECKO^*mice after I/R injury. We next considered *Ripk3*-deficient ECs as a potential source of IL-6 and used cultured MS1 cells in which *Ripk3* was deleted with CRISPR/Cas9 (i.e., *Ripk3*-KO). Since TNFα is one of the key inflammatory cytokines secreted during I/R injury^23^, we stimulated wildtype and *Ripk3*-KO MS1 cells with TNFα for 24 hr to simulate injury and measured secreted IL-6 in the supernatant. We found that *Ripk3* deletion led to enhanced IL-6 secretion both at 0 hr and 24 hr after TNFα stimulation (**Figure 2E**), and no differences were seen in other cytokines we measured (**Supplemental Figure S5E-G**). We also found that *Il6* transcript levels were significantly elevated at 4 hr and 8 hr after TNFα stimulation in *Ripk3*-KO MS1 cells over wildtype cells (**Figure 2F**), indicating a role for endothelial RIPK3 in IL-6 production.

### RIPK3 does not regulate NF-κB signaling in MS1 ECs

The NF-κB signaling pathway is a key mediator of inflammation in ECs. TNFα stimulation of ECs activates canonical NF-κB signaling and results in the nuclear translocation of the transcription factors p65/p50 and p50/c-Rel, where they mediate transcription of inflammatory cytokine and adhesion molecule genes^33–35^. Therefore, we assessed p65 phosphorylation and nuclear localization after TNFα stimulation. We found no differences in p65 protein levels (**Supplemental Figure S6A, B**), and although phospho-p65 levels increased at 10, 30, and 60 mins after TNFα stimulation, they did so equivalently in both wildtype and *Ripk3*-KO cells (**Supplemental Figure S6C, D**). Immunocytochemical staining revealed robust p65 nuclear localization after 10, 30, and 60 mins of stimulation, but no obvious differences were detected between wildtype and *Ripk3*-KO cells (**Supplemental Figure S6E**), indicating that canonical NF-κB signaling is not hyperactive in *Ripk3*-KO ECs.

### The IL-6 repressor NRF2 has decreased nuclear localization and transcriptional activity in Ripk3-KO ECs

The transcription factor NRF2 has antioxidant and anti-inflammatory roles in multiple cell types, including ECs^36^. In addition to translocating from the cytoplasm to the nucleus upon activation and promoting the transcription of antioxidant genes during oxidative stress^36–38^, nuclear NRF2 also inhibits *Il6* transcription in macrophages^39^ and ECs^40^. Therefore, we questioned whether NRF2 expression or activity was compromised in RIPK3-deficient ECs. Stimulated NRF2 nuclear translocation can occur between 30 min to 2 hr in ECs^41^, so we assessed NRF2 cytosolic and nuclear expression at baseline or after 1 hr stimulation with TNFα or tert-Butylhydroquinone (tBHQ)—a known activator of NRF2. We found that although cytosolic NRF2 expression remained comparable (**Figure 3A-D**), nuclear NRF2 expression was decreased in *Ripk3*-KO cells both at baseline and after TNFα stimulation (**Figure 3B, C**). When nuclear to cytosolic NRF2 ratios were calculated under all three culture conditions, this value was consistently and significantly decreased in *Ripk3*-KO compared to wildtype MS1 cells (**Figure 3B-D**). We next measured NRF2 gene (*Nfe2l2*) expression and found its transcript levels to be elevated at baseline but decreased after 4 hr of TNFα stimulation in *Ripk3*-KO cells (**Figure 3E**). Despite this increase in NRF2 gene transcripts in baseline *Ripk3*-KO MS1 cells, the NRF2 target genes *Hmox1* (heme oxygenase 1) and *Nqo1* [NAD(P)H quinone dehydrogenase 1] did not have comparably elevated transcripts in these mutant cells (**Figure 3F, G**). However, at 4 hr after TNFα stimulation, *Hmox1* transcripts were significantly decreased, and *Nqo1* transcripts trended downward in *Ripk3*-KO compared to wildtype MS1 cells (**Figure 3F, G**). Together, these data demonstrate that both NRF2 nuclear localization (assessed at 1 hr) and subsequent NRF2 transcriptional activity (assessed at 4 hr) are diminished in RIPK3-deficient ECs after TNFα stimulation. Therefore, diminished anti-inflammatory NRF2 transcriptional activity in ECs may contribute to the increased IL-6 expression we detected in *Ripk3^iECKO^* mice after I/R challenge and in *Ripk3*-KO MS1 cells after TNFα stimulation.

**FIGURE 3.**
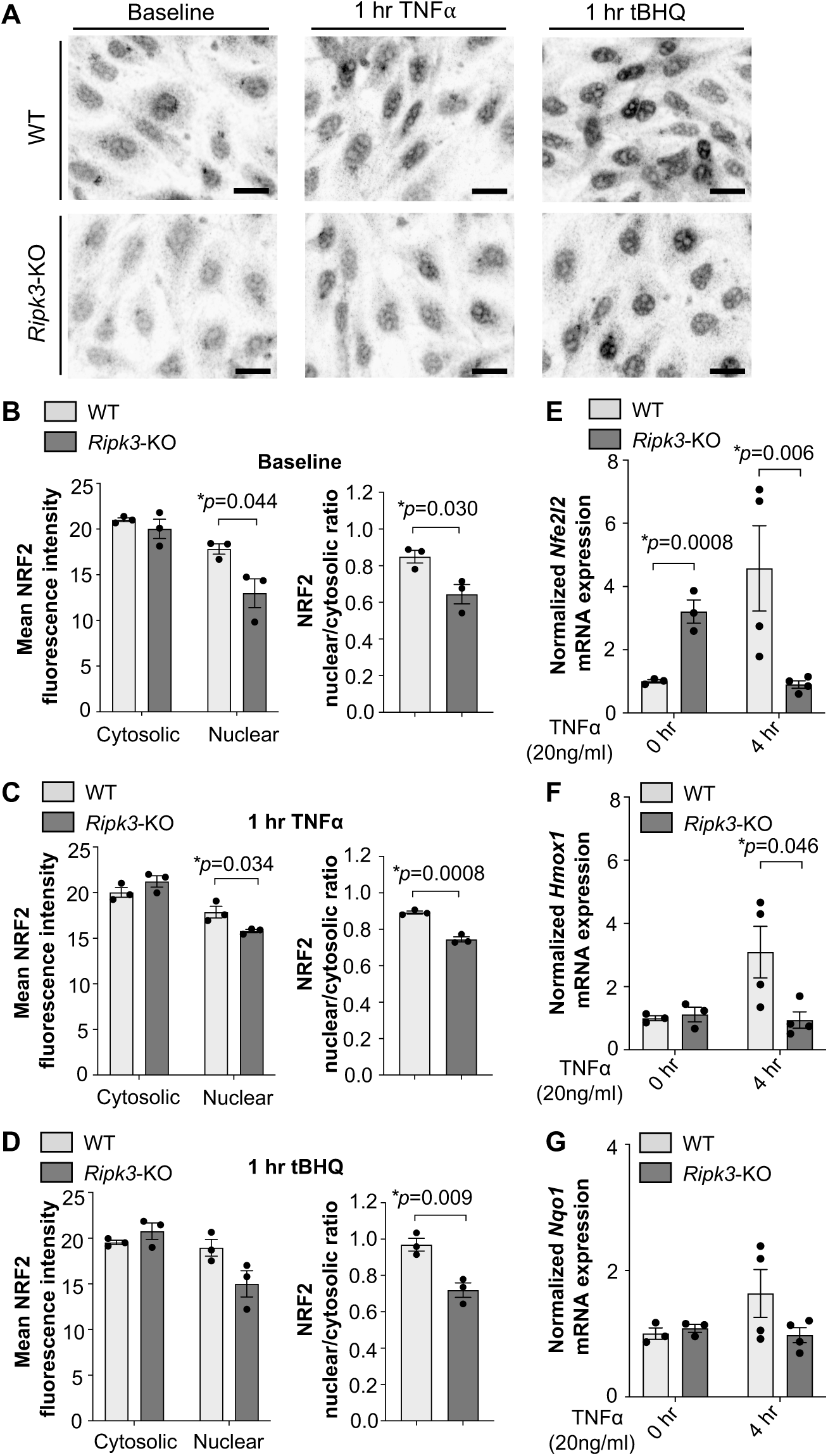
The IL-6 repressor NRF2 has decreased nuclear localization and transcriptional activity in Ripk3-KO endothelial cells (ECs). (**A**) Representative confocal immunofluorescence Z-stack images in grey-scale depicting cytosolic versus nuclear NRF2 expression at baseline and 1 hr following 20ng/mL TNFα or tert-Butylhydroquinone (tBHQ) stimulation in WT and *Ripk3*-KO MS1 ECs. Scale bar: 20 µm. (**B-D**) Mean NRF2 fluorescence intensity in cytosolic and nuclear compartments (left) and calculated NRF2 nuclear to cytosolic ratio (right) in WT and *Ripk3*-KO MS1 ECs at baseline (**B**) or after 1 hr stimulation with TNFα (**C**) or tBHQ (**D**) (n=3/group). ^15^ Transcript expression following stimulation with 20ng/mL TNFα for 0 hr and 4 hr in WT and *Ripk3*-KO MS1 ECs of the following genes: *Nfe2l2* (NRF2) (**E**), *Hmox1* (heme oxygenase-1) (**F**), and *Nqo1* (**G**) (n=3-4/group). *P* values were determined by unpaired t-tests for each treatment condition between the two groups (**B**-**G**), and data that failed an equal variance F-test were log-transformed before analysis (**E**). \**P* <0.05 *vs*. Control. Summary data are the mean ± SE.

### Endothelial adhesion molecules are upregulated in vivo and in vitro after EC-specific deletion of Ripk3

Because immune cell recruitment via endothelial adhesion molecules and extravasation between ECs is another common source of vascular permeability during an inflammatory response^42–46^, we next assessed the expression of VCAM-1 and ICAM-1 in various organs from *Ripk3^iECKO^* mice. Tissue immunoblots revealed that VCAM-1 protein levels were elevated in the small intestines of *Ripk3^iECKO^* mice compared to *Ripk3^WT^* mice at baseline (**Figure 4A**). However, a similar elevation in VCAM-1 was not observed in baseline *Ripk3^iECKO^* lungs, livers, and kidneys (**Figure 4B**), which underscores the emerging biology of organotypically differentiated vascular beds^47^ and hints at how *Ripk3* deletion leads to organ-specific vascular phenotypes after I/R injury. We also stimulated wildtype and *Ripk3*-KO MS1 cells with TNFα and found that VCAM-1 protein expression was increased in *Ripk3*-KO cells at 24 hr post-stimulation compared to wildtype cells by immunoblotting (**Figure 4C**). We saw a similar elevation in VCAM-1 expression when we performed immunocytochemistry on wildtype and *Ripk3*-KO MS1 cells stimulated with TNFα, particularly at EC junctions marked by VE-cadherin (**Supplemental Figure S7A, white arrows**). To determine whether *Vcam1* transcripts were also elevated in stimulated *Ripk3*-KO MS1 cells, we performed quantitative reverse transcription PCR at 4 and 8 hr since transcription increases in the first few hours after TNFα treatment and precedes protein expression. However, we found no difference in *Vcam1* expression between wildtype and *Ripk3*-KO cells (**Figure 4D**).

**FIGURE 4.**
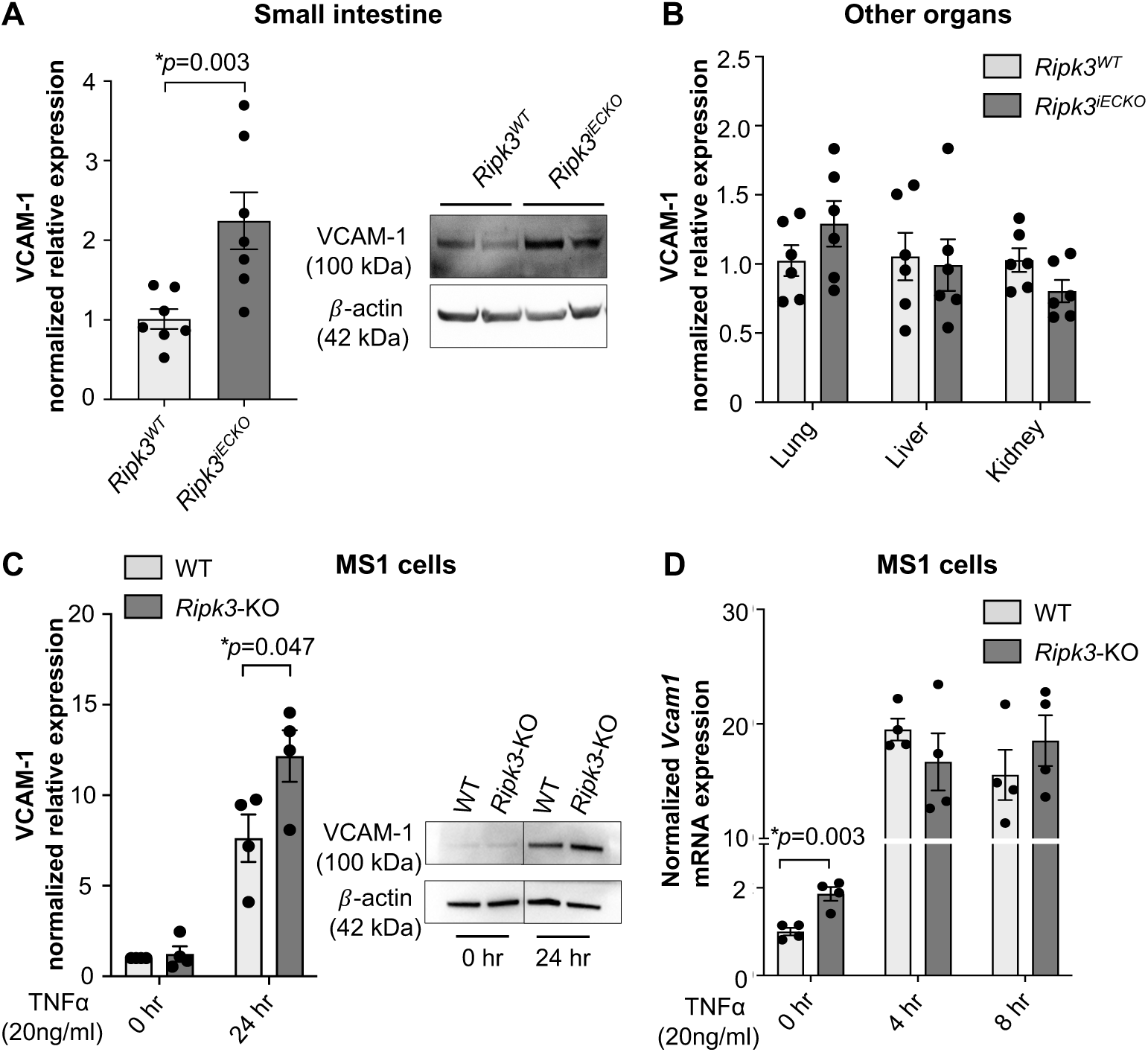
Vascular cell adhesion molecule-1 (VCAM-1) is upregulated in vivo and in vitro after endothelial cell (EC)-specific deletion of Ripk3. (**A**) VCAM-1 expression in the small intestine assessed by immunoblot in baseline *Ripk3^WT^* and *Ripk3^iECKO^* mice (n=7/group). (**B**) VCAM-1 expression in the indicated organs assessed by immunoblot in baseline *Ripk3^WT^* and *Ripk3^iECKO^* mice (n=6/group). (**C**) VCAM-1 expression assessed by immunoblot in WT and *Ripk3*-KO MS1 ECs stimulated with 20ng/ml TNFα for 24 hr (n=4/group). (D) Vcam1 transcript expression in WT and *Ripk3*-KO MS1 ECs following stimulation with 20ng/mL TNFα for 0 hr, 4 hr and 8 hr (n=4/group). *P* values were determined by unpaired t-tests for each treatment condition between the two groups (**A**-**D**), and data that failed an equal variance F-test were log-transformed before analysis (**C**). \**P* <0.05 *vs*. Control. Summary data are the mean ± SE.

When we performed similar analyses of baseline ICAM-1 expression in *Ripk3^WT^* and *Ripk3^iECKO^* mice, tissue immunoblots revealed a significant elevation of ICAM-1 protein expression in mutant lungs (**Figure 5A**). However, ICAM-1 levels were comparable in the small intestine, liver, and kidney between *Ripk3^WT^* and *Ripk3^iECKO^* mice (**Figure 5B**). In cultured MS1 cells, we saw an increase in ICAM-1 levels in *Ripk3*-KO cells at 24 hr after TNFα stimulation (**Figure 5C**). As with VCAM-1, this elevated ICAM-1 in *Ripk3*-KO cells was notable at EC junctions by immunocytochemistry (**Supplemental Figure S7B, white arrows**). However, no differences in *Icam1* transcripts were seen in wildtype and *Ripk3*-KO MS1 cells at 4 and 8 hr after TNFα treatment (**Figure 5D**). Altogether, these data indicate that endothelial RIPK3 prevents excessive VCAM-1 and ICAM-1 protein expression in specific organs after an inflammatory stimulus and likely does so through a post-transcriptional mechanism.

**FIGURE 5.**
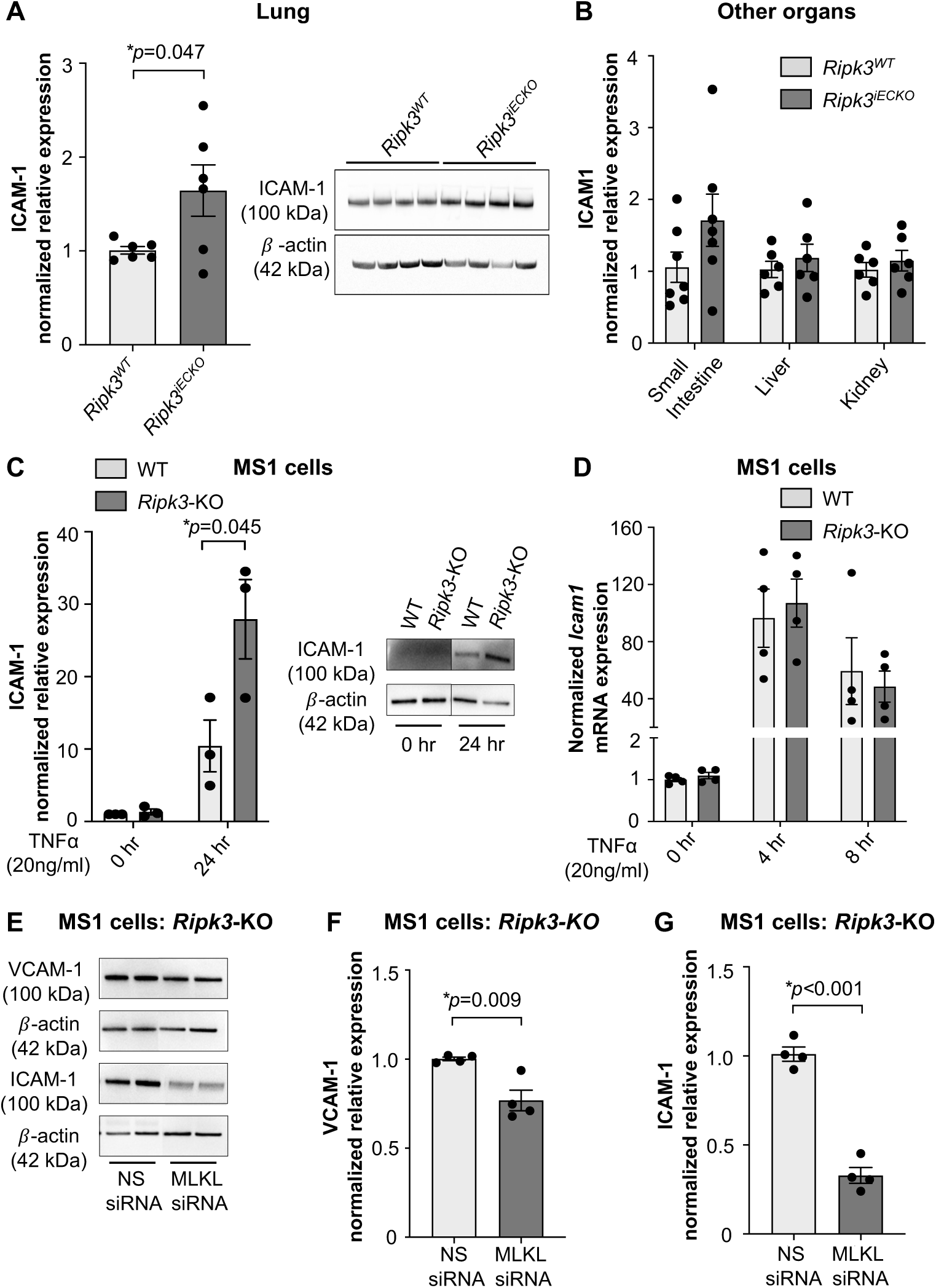
Intercellular adhesion molecule-1 (ICAM-1) is upregulated in vivo and in vitro after endothelial cell (EC)-specific deletion of Ripk3. (**A**) ICAM-1 expression in the lung assessed by immunoblot in baseline *Ripk3^WT^* and *Ripk3^iECKO^* mice (n=7/group). (**B**) ICAM-1 expression in the indicated organs assessed by immunoblot in baseline *Ripk3^WT^* and *Ripk3^iECKO^*mice (n=6/group). (**C**) ICAM-1 expression assessed by immunoblot in WT and *Ripk3*-KO MS1 ECs stimulated with 20ng/mL TNFα for 24 hr (n=4/group). (**D**) *Icam1* transcript expression in WT and *Ripk3*-KO MS1 ECs following stimulation with 20ng/mL TNFα for 0 hr, 4 hr and 8 hr (n=4/group). (**E**) VCAM-1 and ICAM-1 immunoblots in *Ripk3*-KO MS1 ECs after treatment with nonspecific (NS) or MLKL siRNAs and 24 hr of subsequent TNFα stimulation (20ng/mL). (**F**) VCAM-1 expression in *Ripk3*-KO cells following MLKL knockdown and TNFα stimulation, as in E. (**G**) ICAM-1 expression in *Ripk3*-KO cells following MLKL knockdown and TNFα stimulation, as in E. *P* values were determined by unpaired t-tests for each treatment condition between the two groups (**A**, **B**, **C**, **D**, **F** and **G**), and data that failed an equal variance F-test were log-transformed before analysis (C). \**P* <0.05 *vs*. Control. Summary data are the mean ± SE.

MLKL—which is phosphorylated by RIPK3 during necroptotic cell death—has been reported instead to promote post-transcriptional stabilization of adhesion molecules in ECs with very low RIPK3 expression^48^. We therefore sought to determine if MLKL mediated the elevated VCAM-1 and ICAM-1 expression we detected in TNFα-stimulated *Ripk3*-KO cells. We found that MLKL knockdown significantly reduced both VCAM-1 and ICAM-1 protein levels in *Ripk3*-KO cells stimulated for 24 hr with TNFα (**Figure 5E-G**), indicating that MLKL regulates the expression of VCAM-1 and ICAM-1 under inflammatory conditions in RIPK3-deficient ECs.

### Increased numbers of tissue leukocytes are detected in organs of Ripk3^iECKO^ mice following I/R injury

Since I/R injury increases extravasation of leukocytes out of the bloodstream and into tissues^23,31^, we next questioned whether the elevated IL-6 and adhesion molecules we detected in *Ripk3^iECKO^* mice after I/R injury correlated with increased leukocyte recruitment to organs. We first addressed this question *in vitro* by visualizing and quantifying monocyte adhesion to an EC monolayer. We co-incubated fluorescently labeled macrophages (BMA3.1A7) with previously TNFα stimulated wildtype or *Ripk3*-KO MS1 cells for 30 min and found increased macrophage adhesion to *Ripk3*-KO cells compared to wildtype MS1 cells (**Figure 6A**). When fluorescence was quantified in lysed cells with a plate reader, we found that normalized fluorescence was significantly higher in the wells containing *Ripk3*-KO cells (**Figure 6B**). We next immunostained for CD45^+^ leukocytes in the small intestines, livers, lungs, and kidneys of *Ripk3^WT^* and *Ripk3^iECKO^* mice at 24 hr after I/R injury (**Figure 6C**), since these were the organs in which we detected increased vascular permeability in *Ripk3^iECKO^* mice (Figure 1 C-D). We found the largest increase in the numbers of CD45^+^ leukocytes in the small intestines and lungs (**Figure 6D, E**) of *Ripk3^iECKO^* tissues when compared with littermate controls. This corresponded specifically with organs that displayed elevated adhesion molecules in baseline *Ripk3^iECKO^* mice (Figures 4A and 5A).

**FIGURE 6.**
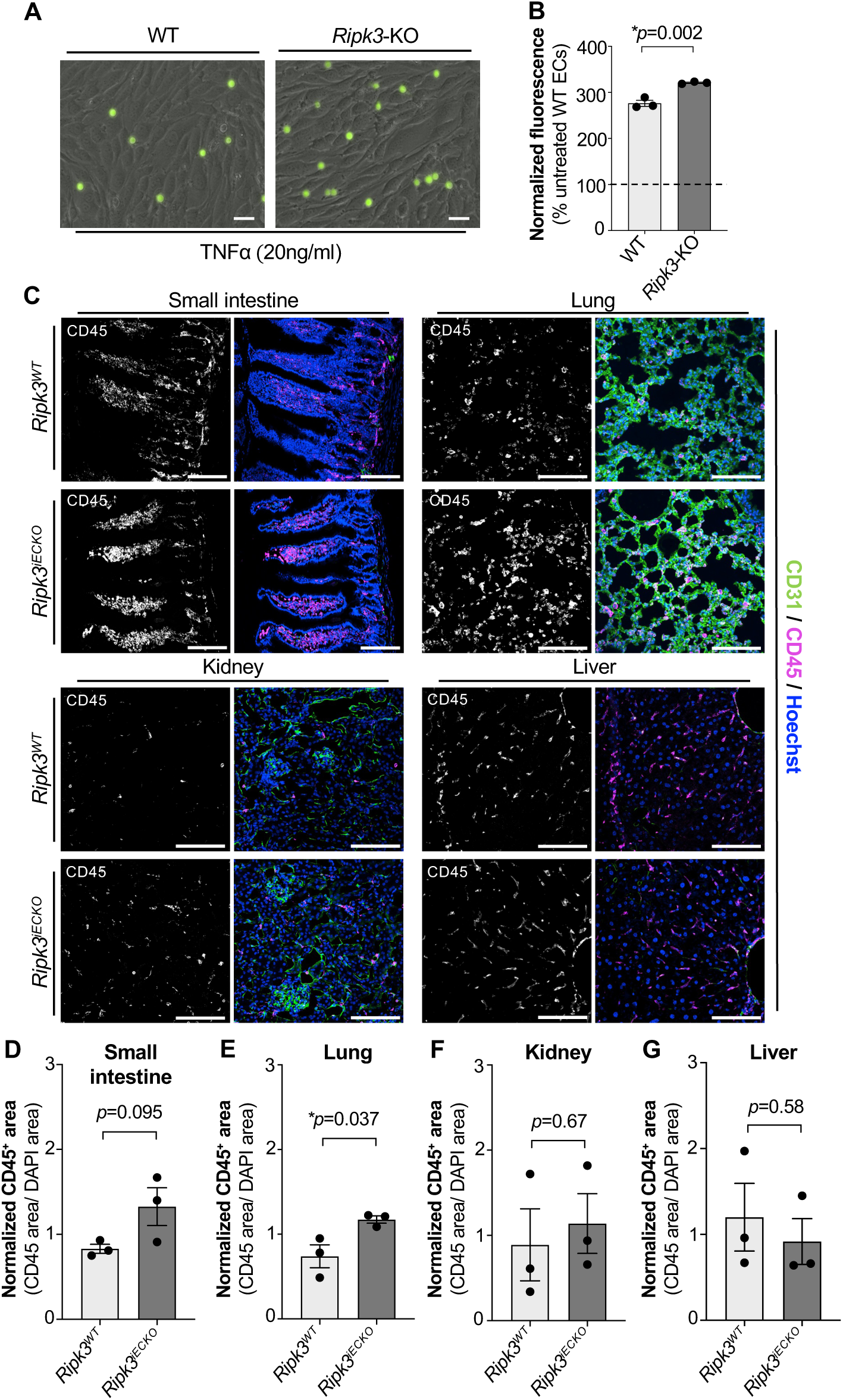
Leukocyte adherence to endothelial cells (ECs) and accumulation in tissues is elevated after EC-specific deletion of Ripk3 under inflammatory conditions. (**A**) Representative images of immortalized BMA3.1A7 macrophages labeled with calcein-AM fluorescent dye (green) and incubated on top of a monolayer of WT and *Ripk3*-KO MS1 ECs that were previously treated with 20ng/mL TNFα for 24 hr. Cells were washed multiple times before imaging. Scale bar: 20 µm. (**B**) Following imaging (as in A), cells were lysed with Triton-X, and fluorescence was measured using a plate reader. Values were normalized to readings from wells in which BMA3.1A7 cells were incubated with untreated WT MS1-ECs (dotted line) (n=3/group). (**C**) CD45^+^ leukocyte accumulation was assessed by immunostaining in various tissues from *Ripk3^WT^* and *Ripk3^iECKO^*mice following 24 hr I/R injury. Scale bar: 100 µm. CD45^+^ cells were quantified from multiple confocal Z-stack images taken from each mouse tissue by normalizing CD45^+^ expression area to total cell nuclear area in small intestines (**D**), lungs (**E**), kidneys (**F**) and livers (**G**) of *Ripk3^WT^* and *Ripk3^iECKO^*mice. Individual data points are averages of 2-4 imaged and quantified sections per mouse (n=3 mice/group). *P* values were determined by unpaired t-tests **(B, D, E, F** and **G**). \**P* <0.05 *vs*. Control. Summary data are the mean ± SE.

### Monocyte reduction in Ripk3^iECKO^ mice rescues small intestinal and lung vascular permeability after I/R injury

Circulating monocytes can compromise the vascular barrier during injury and inflammation via adhesion-mediated signaling and extravasation^44,46,49–51^. We therefore sought to evaluate the contribution of circulating monocytes to vascular hyperpermeability in different organs following I/R injury. PBS- or clodronate-liposome administration to wildtype mice for 48 hr led to complete depletion of F4/80^+^ liver macrophages and partial depletion of F4/80^+^ macrophages in lungs and kidneys but no depletion in the small intestines (**Figure 7A**). Additionally, clodronate-liposome treatment led to 47% depletion of circulating monocytes (**Figure 7B**). Notably, both control and *Ripk3^iECKO^*mice treated with this clodronate-liposome scheme for 48 hr displayed difficulty tolerating isoflurane anesthesia during the ischemic period of our I/R injury surgical procedure; 1-2 mice from each group died during the procedure. Therefore, we continued with the same concentration of clodronate-liposomes rather than increasing the concentration to achieve higher monocyte/macrophage depletion. We performed this clodronate-liposome administration scheme in *Ripk3^WT^* and *Ripk3^iECKO^* mice followed 48 hr later by I/R injury surgeries and subsequent EBD leakage assays to measure vascular permeability (**Figure 7C**). We found that the elevated vascular permeability previously detected in the small intestinal regions of *Ripk3^iECKO^* mice (Figure 1C) was rescued after monocyte reduction (**Figure 7D**). Likewise, the elevated permeability we had seen in the lungs of *Ripk3^iECKO^* mice (Figure 1D) was rescued after clodronate-liposome administration (**Figure 7E**). However, vascular permeability in *Ripk3^iECKO^* livers and kidneys was not rescued as impressively after monocyte reduction (**Figure 7E**). Notably, the small intestines and lungs were the two organs that displayed elevated adhesion molecules in baseline *Ripk3^iECKO^* mice (Figures 4A and 5A) and the most inflammation after I/R injury (Figures 6C-G). Altogether, these data suggest that circulating monocytes are recruited to organs with elevated endothelial adhesion molecules in *Ripk3^iECKO^* mice and contribute to vascular permeability in these organs after I/R injury.

**FIGURE 7.**
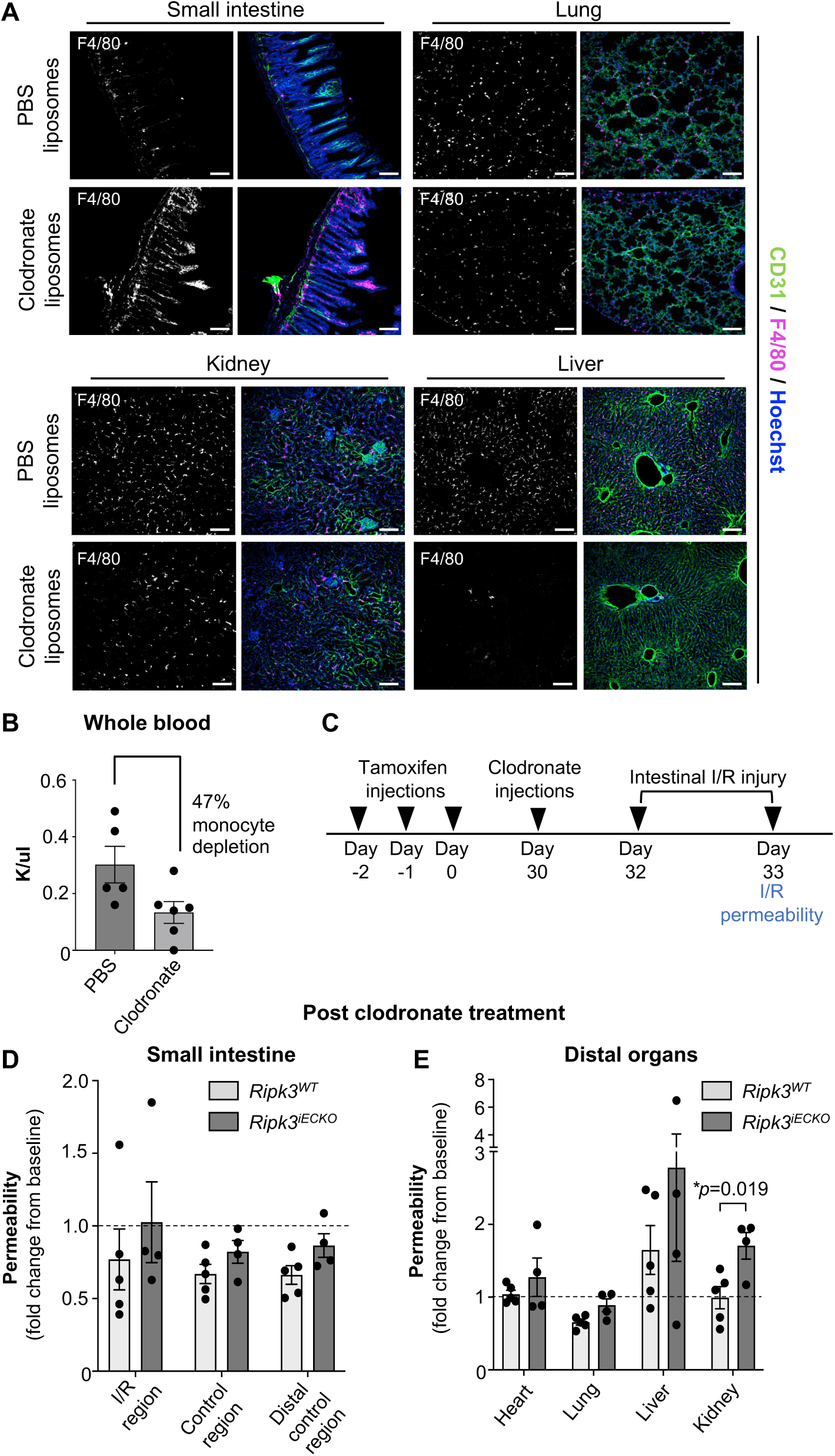
In vivo monocyte depletion rescues small intestinal and lung vascular permeability after intestinal I/R injury in Ripk3^iECKO^ mice. (**A**) Representative confocal Z-stack images of F4/80^+^ macrophage depletion in indicated organs 48 hr following administration of PBS liposomes or clodronate liposomes. Scale bar: 100 µm. (**B**) Circulating blood monocyte depletion using PBS liposomes versus clodronate liposomes (n=5/group). (**C**) Timeline for detecting vascular permeability after clodronate-induced monocyte depletion and intestinal I/R injury; tamoxifen will be administered at 8 weeks of age. (**D**) Small intestinal vascular permeability after 24 hr intestinal I/R injury in the injury region (I/R) and control regions (control, distal control) following clodronate-induced monocyte depletion (n=4-5/group). Fold change is calculated from baseline permeability (see Figure 1B, dotted line). (**E**) Vascular permeability in the indicated organs after 24 hr intestinal I/R injury following clodronate induced monocyte depletion (n=4-5/group). Fold change is calculated from baseline permeability (see Figure 1B, dotted line). *P* values were determined by unpaired t tests for each treatment condition between the two groups (**D** and **E**), and data that failed an equal variance F-test were log-transformed before analysis (**D**). \**P* <0.05 *vs*. Control. Summary data are the mean ± SE.

## DISCUSSION

Our current study addresses the role of endothelial RIPK3 in intestinal I/R injury-induced vascular permeability. Based on previous work by our lab and others^5,13,14,52,53^, we predicted that RIPK3 would be elevated in ECs after ischemic injury and would subsequently promote vascular permeability. While we did find RIPK3 elevated in mesenteric vessels exposed to ischemia and I/R, we were surprised to find vascular permeability increased after I/R injury in several organs in *Ripk3^iECKO^* mice compared to littermate controls. This increase in vascular permeability correlated with (1) elevated IL-6 in serum and small intestinal tissue, (2) enhanced expression of the endothelial adhesion molecules ICAM-1 in lungs and VCAM-1 in small intestines, and (3) predominant immune cell infiltration into lungs and small intestines. Altogether, our data demonstrate that endothelial RIPK3 plays a beneficial role in maintaining vascular integrity after I/R injury, which contradicts the detrimental reputation that is often associated with RIPK3 due to its contributions to programmed cell death pathways and inflammasome activation in pathological contexts^54^. Notably, this is not the first time we have found beneficial roles for endothelial RIPK3. We previously reported that RIPK3 promotes developmental angiogenesis in the embryonic hindbrain and newborn retina by regulating growth factor receptor trafficking^29^. We also reported that apolipoproteindeficient (*Apoe^-/-^*) *Ripk3^iECKO^* mice fed a high fat diet develop more atherosclerotic lesions in their descending aortas than littermate controls^28^. Importantly, this increased atherosclerosis correlated with elevated expression of the adhesion molecule E-selectin in cultured RIPK3-knockdown ECs. Therefore, our previous atherosclerosis study correlates with the current study in terms of revealing protective roles for endothelial RIPK3 in inflammatory challenge models and with intriguing similarities in the regulation of adhesion molecules that mediate immune cell extravasation.

Although we saw vascular permeability elevated in *Ripk3^iECKO^* mice after I/R injury, it is worth noting that unchallenged *Ripk3^iECKO^* mice displayed comparable or even decreased vascular permeability compared to control mice in all the organs we assessed. This may reflect innate organotypic differences in vascular beds, including baseline expression of RIPK3 targets that modulate vascular permeability. Regardless, the decreased vascular permeability we saw in the lungs and kidneys of unchallenged *Ripk3^iECKO^* mice suggests that RIPK3 antagonizes endothelial barrier function in these homeostatic organs via mechanisms that will require further investigation. Additionally, RIPK3 behaves differently when regulating vascular permeability in baseline versus stimulated ECs, which raises questions about how the complicated environment of the I/R challenge model triggers RIPK3 to assume its protective role. Future studies should explore contributions of hypoxia, reactive oxygen species, nitric oxide, and cytokine signaling toward this shift in endothelial RIPK3 activity in vivo. Another surprise that emerged from our study was the finding that RIPK3 suppresses vascular permeability in an organotypic fashion after I/R challenge. Not only was permeability elevated in regions of the small intestine downstream of unclamped mesenteric vessels in *Ripk3^iECKO^*mice compared to control littermates, but it was also increased in the lung, liver, and kidneys of mutant mice after I/R injury. However, no change in vascular permeability was seen in the hearts of *Ripk3^iECKO^* mice after I/R injury. The cause of this organ-specific variability in the RIPK3-mediated vascular permeability response to I/R injury is unclear, but it likely involves complex reactions to inflammatory cytokines and endothelial adhesion molecules, as discussed further below.

Inflammatory cytokines are known contributors to vascular permeability after I/R injury^23,31,32^, and our current study reveals that IL-6 is significantly elevated after I/R injury in the serum and small intestinal tissue of *Ripk3^iECKO^* mice. We also found that IL-6 protein and transcripts were elevated in *Ripk3*-KO MS1 cells after stimulation with TNFα. Although ECs are a known source of IL-6 under inflammatory conditions^55^, our data are the first to indicate that RIPK3 serves as a brake on endothelial IL-6 production. Mechanistically, we believe that RIPK3 regulates IL-6 expression via the transcription factor NRF2, which is activated by oxidative stress and plays antiinflammatory roles in several different cell types, including ECs^36–40^. Notably, NRF2 can inhibit *Il6* transcription indirectly by upregulating the gene encoding the anti-oxidant *Hmox1*^37^ or directly by binding near the transcription start site of *Il6* and inhibiting RNA polymerase II recruitment^39,40^. We found impaired NRF2 nuclear localization and transcriptional activity when *Ripk3*-KO MS1 cells were stimulated with TNFα. Therefore, RIPK3 may suppress excessive IL-6 secretion from ECs under inflammatory conditions by promoting NRF2 activation, nuclear transport, and *Il6* transcriptional inhibition. Because phosphorylation of different sites within NRF2 can positively or negatively regulate its stability and intracellular localization^56^, it is possible that RIPK3 kinase activity directly or indirectly impacts NRF2 activity. Regardless, our work suggests that the regulatory relationship between endothelial RIPK3 and NRF2 is a rich area for future investigation.

IL-6 uses two mechanisms to mediate its biological effects. Classical IL-6 signaling occurs when the cytokine binds to a cell surface complex consisting of the IL-6 receptor α (IL-6Rα) and the ubiquitously expressed 130-kilodalton signal-transducing β-receptor subunit (gp130). Cells that do not express IL-6Rα—including ECs—undergo IL-6 signaling when the cytokine binds a soluble form of IL-6Rα, and this complex subsequently interacts with membrane bound gp130 to mediate trans-signaling^57^. Classical IL-6 signaling typically regulates homeostatic processes, while IL-6 trans-signaling occurs in cardiovascular diseases and mediates inflammation^57^. This pro-inflammatory trans-signaling also promotes vascular permeability through activation of intracellular JAK/STAT3 proteins^55,58–61^. Therefore, we believe that the elevated IL-6 we detected in the serum of *Ripk3^iECKO^*mice contributed to increased I/R-induced vascular permeability through trans-signaling on ECs. We also suspect that ECs are a major source of the elevated IL-6 we found in *Ripk3^iECKO^* mice after I/R injury, which would be consistent with our *in vitro* data. Although immune cells are usually the principal sources of this cytokine in inflammatory pathologies^62^, we found no evidence of *Ripk3* deletion in immune cells from *Ripk3^iECKO^* mice. Likewise, whole blood removed from the mutant mice and stimulated *ex vivo* did not show elevated IL-6 secretion compared to blood from control mice. A genetic model in which *Il6* is selectively reduced in ECs would help clarify the contribution of these cells to the elevated IL-6 in I/R injury-challenged *Ripk3^iECKO^* mice. Regardless of whether ECs or tissue resident cells are the major source of the elevated IL-6 in I/R-challenged *Ripk3^iECKO^* mice, it is important to note that IL-6 is garnering significant attention for its ability to promote cytokine storms, particularly in the context of severe infection (e.g., influenza virus), acute respiratory distress syndrome associated with COVID-19, and Car-T cell therapy^55,63^. Therefore, it will be interesting to learn in the future whether endothelial RIPK3 regulates IL-6 production in these and additional inflammatory settings.

Another novel finding from the current study is that the endothelial adhesion molecules VCAM-1 and ICAM-1 were elevated in unchallenged *Ripk3^iECKO^* mice in a tissue-specific manner. VCAM-1 was selectively upregulated in the small intestine, and ICAM-1 was upregulated in the lungs of *Ripk3^iECKO^* mice compared to littermate controls, but these adhesion molecules were not elevated in the other tissues we analyzed that displayed increased vascular permeability after I/R injury (i.e., kidney and liver). The fact that these adhesion molecules were upregulated prior to I/R injury indicates that RIPK3 plays a direct role in regulating their expression rather than an indirect role through elevated IL-6 signaling that occurs with injury. Because we saw VCAM-1 and ICAM-1 proteins upregulated without an elevation in their corresponding transcripts in *Ripk3*-KO MS1 cells after TNFα challenge, we were drawn to a recent report by Dai et al that MLKL can interact with the RNA binding protein RBM6 to stabilize transcripts of VCAM-1, ICAM-1, and E-selectin in ECs with low levels of RIPK3 and promote the cell surface expression of these adhesion molecules^48^. This study also showed that MLKL-mediated adhesion molecule expression facilitates leukocyte binding to vascular walls and extravasation in an inflammatory challenge model. While MLKL is commonly known as a target of RIPK3 kinase activity and an effector of cellular necroptosis, the Dai et al report suggests that MLKL can assume other important cellular roles in ECs lacking RIPK3. With this in mind, we assessed whether MLKL might mediate the post-transcriptional VCAM-1 and ICAM-1 upregulation we saw in TNFα-treated *Ripk3*-KO MS1 cells. Indeed, we found that MLKL-knockdown suppressed the elevation of these adhesion molecules in the mutant cells, which indicates that MLKL may also mediate VCAM-1 and ICAM-1 upregulation in *Ripk3^iECKO^*mice. Future work could address whether organ-specific differences in endothelial MLKL or RBM6 expression might explain the respective small intestinal- and lung-specific upregulation we saw in VCAM-1 and ICAM-1 expression in *Ripk3^iECKO^* mice. For now, we can emphasize that a consistent theme of pathological adhesion molecule elevation is emerging from our analyses of endothelial *Ripk3* knockout mice, based on our previous atherosclerosis study^28^ and the current I/R challenge model.

Because VCAM-1 and ICAM-1 serve as docking sites for circulating leukocytes that can subsequently extravasate out of the blood stream^63–65^, the regulation of adhesion molecule expression is highly relevant to tissue inflammation. Moreover, leukocyte binding to adhesion molecules induces intracellular signaling that compromises EC barrier function and promotes vascular permeability^44,46,49–51^. We found that macrophages adhered more robustly to cultured *Ripk3*-KO MS1 cells than to wildtype cells, and our in vivo immunostaining indicated that CD45^+^ leukocytes were more abundant following I/R injury in *Ripk3^iECKO^*tissues than in those from control littermates. Considering these findings, we were particularly interested to find that clodronate liposomes significantly rescued I/R-induced vascular permeability in *Ripk3^iECKO^* lungs and small intestines—the two tissues in which adhesion molecules were selectively upregulated in baseline mutant mice. Notably, resident macrophage depletion was not impressive in these tissues with our clodronate liposome administration scheme. Therefore, we believe that circulating monocytes—which were depleted by ∼47% with clodronate liposomes—were more impactful contributors to permeability than were tissue resident macrophages in these organs after I/R challenge. Although we saw robust depletion of resident liver macrophages with clodronate liposomes, this did not impact hepatic vascular permeability after I/R challenge in *Ripk3^iECKO^*mice. Altogether, these data indicate that circulating monocytes impact vascular permeability after I/R challenge—potentially through a combination of their adhesion, extravasation, and local cytokine release. Furthermore, we speculate that organs such as the liver and kidney, which did not have elevated adhesion molecule expression and did not have vascular permeability rescued by clodronate treatment, were more impacted by increased IL-6 secretion in *Ripk3^iECKO^*mice.

Altogether, this study highlights a beneficial role for endothelial RIPK3 in the complex pathophysiological response to intestinal I/R injury. It also demonstrates that RIPK3 regulates organotypic vascular adhesion molecule expression and endothelial IL-6 secretion, which may have implications for multiple inflammatory conditions. These findings can be added to the growing list of non-necroptotic functions for RIPK3 in ECs and should raise awareness that the use of pharmacological necroptosis inhibitors for treating ischemic diseases could have unintended effects on the vasculature.

## ONLINE METHODS

### Mouse lines and treatment

All mouse studies were performed in accordance with the NIH Guide for the Care and Use of Laboratory Animals and were approved by the Institutional Animal Care and Use Committee at the Oklahoma Medical Research Foundation (protocol # 23-12). The mice used in this study were maintained on a C57Bl/6J background and were housed in a single room of the animal facility with a 12-hour light-dark cycle; food and water were provided *ad libitum*. Both female and male adult mice were used in all experiments (see **Supplemental Table S1**). *Ripk3*-floxed mice^5^, *Cdh5(PAC)-Cre^ERT2^*mice (gift of Ralf Adams, Max Planck Institute for Molecular Biomedicine; Taconic; #13073)^66^, RIPK3-GFP mice (*Ripk3-gfp^fl/fl^*; The Jackson Laboratory; #030284)^27^, and *ROSA^mT/mG^* mice (*Gt(ROSA)26Sor^tm4(ACTB-tdTomato,-EGFP)Lou^/J*; The Jackson Laboratory; #007576;)^30^ were previously described. *Ripk3^fl/fl^* mice were crossed with *Cdh5(PAC)-Cre^ERT2^*mice to generate *Ripk3^fl/fl^;iCdh5(PAC)-Cre^ERT2^* mice (i.e., *Ripk3^iECKO^*), and *Ripk3^fl/fl^* littermates were used as controls (i.e., *Ripk3^WT^*). Tamoxifen (Sigma; #T5648) dissolved in peanut oil (10mg/ml stock concentration) was administered intraperitoneally (i.p.) at 2mg/day/mouse for 3 consecutive days in 8-week-old mice; experiments were conducted 4 weeks later to allow for full gene deletion. A similar tamoxifen administration scheme was used to assess Cre activity in *ROS-A^mT/mG^;iCdh5(PAC)-Cre^ERT2^* mice. Genotyping for each allele was performed using primers listed in **Supplemental Table S2**.

### Intestinal ischemia-reperfusion (I/R) injury

After mice were anesthetized using isoflurane (3%) and extended-release buprenorphine (1.0 mg/kg dose; ZooPharm; #1Z-74000-222510) was administered as an analgesic, a mid-line 3 cm laparotomy was performed, and the cecum and ileum were isolated. As described previously^67^, atraumatic microvascular clips (Roboz; #RS-5420) were used to occlude first-order branches of the superior mesenteric artery feeding the ileum. Ischemia was maintained for 1 hr using 1-1.5% isoflurane anesthesia, and body temperature was maintained at 37°C using a water circulating heating pad. Tissues were kept moist by irrigation with sterile saline pre-warmed to 37°C throughout the 1 hr period. At the end of the ischemic period, the microvascular clips were removed, the small intestines were placed back in the abdominal cavity, and the abdominal wall was sutured. Mice were then placed in a clean cage for a 24 hr reperfusion period before euthanasia via carbon dioxide inhalation at a flow rate of 3 liter/min followed by cervical dislocation. Unclamped adjacent vessels and their downstream small intestinal segments were considered as internal control regions. In a separate cohort of mice, sham I/R injuries were performed as described above without the occlusion of first-order branches of the superior mesenteric artery feeding the ileum.

### Mesenteric arterial en face preparations and immunolabeling for endothelial RIPK3 expression

Primary and secondary mesenteric arterial branches of the superior mesenteric artery exposed to either 1 hr ischemia or 24 hr I/R injury along with adjacent control (unclamped) mesenteric arterial branches were isolated from *Ripk3-gfp^fl/fl^*mice. Vessels were then carefully cut open longitudinally and pinned to a sylgard-coated 12-well plate for subsequent immunolabeling of GFP/RIPK3 in mesenteric arterial endothelial cells^68^.

### In vivo vascular permeability

Vascular permeability was measured using an Evans blue dye (EBD) assay. Mice were first anesthetized using isoflurane (3%). 1% EBD (Sigma; #E2129) was administered retro-orbitally (4 µL/g bodyweight) and allowed to circulate for 30 mins. Afterward, mice were euthanized via carbon dioxide inhalation at a flow rate of 3 liter/min followed by cervical dislocation, and organs (small intestine, liver, lung, kidney, and heart) were harvested. Organ wet weights were measured, then organs were dried for 24-48 hours at 37°C before dry weights were measured. Organs were then incubated in 500 µL formamide for 48 hr at 56°C to extract EBD from the tissues. Extravasated EBD was measured spectrophotometrically at 620 nm and 740 nm, and values were corrected for hemoglobin using the following formula: OD_620_ - (1.426 X OD_740_ + 0.03), as previously described^69^. The concentration of EBD extravasated from specific tissues was calculated using a standard curve of known EBD concentrations, and final values were normalized to organ dry weight. 70 kDa fluorescein isothiocyanate-dextran (FITC-dextran; Sigma; #46945) was also used to measure vascular permeability. FITC-Dextran was injected retro-orbitally at a final dose of 100mg/kg bodyweight following a 24 hr I/R injury challenge. FITC-dextran was allowed to circulate for 30 mins, and mouse tissues were perfusion-fixed with 4% paraformaldehyde. Organs were then collected and cryoembedded for sectioning and fluorescent microscopy.

### In vivo circulating monocyte depletion

Monocyte depletion from circulating blood was first calculated in a wildtype cohort of mice injected i.p. with either PBS-liposomes or clodronate-liposomes (Encapsula Nano Sciences; #CLD-8901)^70^ at 100 µL/10 g body weight. Liposomes were allowed to circulate for 48 hr before mice were euthanized via carbon dioxide inhalation at a flow rate of 3 liter/min followed by cervical dislocation and blood was collected by cardiac puncture. A hemavet (Drew Scientific; #HV03586) was used to measure circulating monocyte numbers in the blood. Subsequently, monocyte depletion was performed in littermate control and *Ripk3^iECKO^* mice, and I/R injury was performed as described above after clodronate liposomes had circulated for 48 hr.

### Endothelial cell (EC) and blood leukocyte isolation from mice

To isolate ECs, mouse lungs were enzymatically dissociated with collagenase B (Roche; #34324621, 1mg/ml) and dispase II (Sigma; #D4693, 1mg/ml) for 50 min at 37°C. DNase I (Ambion; #AM2224) was added to the collagenase/dispase mix to remove genomic DNA. Next, sheep anti-rat Dynabeads (Invitrogen; #11035) coated with anti-CD31 antibody (BD Pharmingen; MEC13.3, #553370) were incubated with the lung single-cell suspensions for 45 min at 4°C. ECs were then magnetically separated from the rest of the lung cells, and both EC and non-EC cell samples were placed in TRIzol Reagent for RNA isolation. To isolate peripheral blood leukocytes, mice were euthanized via carbon dioxide inhalation at a flow rate of 3 liter/min followed by cervical dislocation, and blood was then collected by cardiac puncture with a syringe pre-filled with 100 µl of 3.2% sodium citrate and placed in 10 ml ACK lysis buffer (Gibco; #A1049201) for 15 min at 25°C for erythrocyte lysis. Blood samples were then centrifuged, supernatant was aspirated, and cells were suspended in RNase free PBS. Samples were centrifuged again to remove the PBS before adding TRIzol Reagent for RNA isolation.

### EC culture

The MS1 adult murine pancreatic EC line (ATCC; #CRL 2279) was cultured in Dulbecco’s Modified Eagle Medium (DMEM) (Gibco; #11960-044) supplemented with 10% fetal bovine serum (FBS) (Fisher Scientific; 3SH30910.03) and 1% antibiotics (streptomycin/amphotericin; Sigma; #A5955). *Ripk3*-KO MS1 cells, which were generated by CRISPR-Cas9 modification^71^, were cultured identically. TNFα (R&D Systems; #410-MT-010/CF) was administered at 20 ng/mL for designated times. For oxygen glucose deprivation (OGD) studies, DMEM was replaced with no glucose DMEM (Gibco; #11966025) before MS1 cells were placed in normoxia (19% O2) or hypoxia (1% O2) chambers (BioSpherix) for 6 hr, 8 hr or 12 hr. For NRF2 studies, cells were incubated with 20 ng/mL TNFα or 50 µM tert-Butylhydroquinone (tBHQ; Sigma; #112941) for 1 hr before immunostaining for NRF2. For siRNA-mediated knockdown of MLKL in *Ripk3*-KO MS1 cells, nonspecific (NS) control siRNA (Thermo; #4390844) or MLKL siRNA (Thermo; #4390771) was transfected with Lipofectamine RNAiMAX reagent (Invitrogen; #56532) for 24 hr before stimulating cells with TNFα for an additional 24 hr.

### Monocyte adhesion assay

The BMA3.1A7 immortalized mouse macrophage line (Applied Biological Materials; #T06730) was cultured in RPMI 1640 medium (Sigma; #R8758) supplemented with 10% FBS (Fisher Scientific; 3SH30910.03) and 1% antibiotics (streptomycin/amphotericin; Sigma; #A5955). BMA3.1A7 cells were labeled with 4µM Calcein AM (AAT Bioquest; #22004), as previously described^48,72^. They were then incubated for 30 min at 37°C on top of a monolayer of MS1 ECs (ATCC; #CRL 2279) that had been pretreated with 20 ng/mL TNFα (R&D Systems; #410-MT-010/CF) for 24 hr. Nonadherent BMA3.1A7 macrophages were removed by multiple washing steps with Hanks’ Balanced Salt Solution (HBSS; Gibco; #14175) before remaining cells were imaged using a Nikon TiE Eclipse epifluorescence microscope. Cells were then lysed in 1% Triton X-100 (Thermo Fisher Scientific; #A16046.AE) for 10 min, and fluorescence was measured at excitation wavelength 490 nm and emission wavelength 525 nm and normalized to a control group in which MS1 ECs were not incubated with macrophages.

### Ex vivo whole blood cytokine release assay

Cytokines in whole blood were measured using a cytokine release assay, as previously described^73–75^. Mice were first euthanized via carbon dioxide inhalation at a flow rate of 3 liter/min followed by cervical dislocation, and blood was then collected through a cardiac puncture into tubes with the anticoagulant lepirudin (50 µg/mL; gifted by Dr. Florea Lupu) to prevent blood clotting. Collected blood was then incubated with 1 µg/mL bacterial lipopolysaccharide (LPS; Sigma; #L3012) or PBS for 2 hr and 4 hr. Plasma samples were collected at the end of each incubation period for downstream cytokine measurements using a multiplex immunoassay.

### In vivo and in vitro cytokine measurements

Serum cytokines in mice were measured following a 4 hr I/R injury. Blood leukocyte cytokine release in plasma was measured after stimulating leukocytes with LPS, as described above. MS1 cells were stimulated with 20 ng/mL TNFα for 24 hr, and secreted cytokines were measured in the cell-free supernatant. Serum, plasma, or supernatant cytokines were measured using a custom ProcartaPlex^TM^ Multiplex Immunoassay (Thermo Fisher). The plates were read on a Bio-Plex 200 array reader (Bio-Rad). Samples were run in duplicate, and cytokine levels were quantified by comparing to a 4PL algorithm standard curve specific to each analyte^76^.

### Immunofluorescence

Organs were collected from euthanized mice and briefly rinsed in PBS before fixing overnight with 4% paraformaldehyde at 4°C. Organs were then incubated in 10%, 15%, and 20% sucrose (1 hr each) and placed in a 1:1 mixture of 20% sucrose and optical cutting temperature compound (O.C.T; Sakura; #4583) overnight. Organs were then embedded in O.C.T. the next morning in cryomolds, and 20 µm tissue sections were cut with a cryotome (Fisher Scientific; Epredia^TM^ Microm HM525 NX Cryostat). Tissue sections were dried at 37°C, washed in PBS for 5 min to remove O.C.T., permeabilized for 15 min in 0.3% Triton X-100, and blocked in 10% donkey serum (Jackson ImmunoResearch Lab; #102644-006) in PBS for 1 hr. Primary antibodies were diluted in 3% bovine serum albumin (BSA; Rockland; #BSA-50), 0.1% Triton X-100 in PBS and applied to sections overnight at 4°C. Secondary antibodies were diluted in 3% BSA in PBS and applied to sections the next day for 1 hr at room temperature in the dark. Confocal Z-stack images were obtained with a Nikon C2 confocal microscope. Primary antibodies used for immunofluorescence were rabbit-anti-GFP (1:200, Novus Biological; #NB600-303), rat-anti-VE-cadherin (1:100, Abcam; #ab91064), goat-anti-CD45 (1:50, R&D Systems; #AF114), goat-anti-VCAM-1 (1:100, R&D Systems; #AF643), goat-anti-ICAM-1 (1:100, R&D Systems; #AF796), rabbit-anti-NF-κB (nuclear factor kappa-light-chain-enhancer of activated B cells) p65 (1:700, Cell Signaling; #8242), goat-anti-CD31 (1:100, R&D Systems; #AF3628), rat-anti-CD31 (1:100, BD Pharmingen; #553370) and rabbit-anti-NRF2 (1:300, gift from Dr. Scott Plafker at OMRF) ^77^. Secondary antibodies used were Cy5-donkey-anti-rabbit IgG (1:500, Jackson ImmunoResearch), Alexa488-donkey-anti-rat IgG (1:500, Thermo Fisher), and Cy3-donkey-anti-goat IgG (1:500, Jackson ImmunoResearch). Hoechst (20 µg/mL) was added to the secondary antibody incubation^5,78^. A similar staining protocol was used for cultured cells grown on coverslips (Electron Microscopy Sciences; #7229608) and fixed with 4% paraformaldehyde for 15 min.

### Immunoblotting

Frozen mouse tissue samples or cells were first lysed in RIPA buffer (Sigma; #R0278) that included Protease Inhibitor Cocktail (Thermo Fisher; #1860932) and Phosphatase Inhibitor Cocktail (Thermo Fisher; #78420) using a mini bead beater (DENTSPLY; #C321001). Protein concentrations of lysates were then measured with a BCA Protein Assay Kit (Thermo Scientific; #23227) following the manufacturer’s instructions. Samples were diluted in 4X Bolt LDS Sample Buffer (Thermo Scientific; #B0007) with 10X Bolt Sample Reducing Agent (Thermo Scientific; #B0009) and heated at 70°C for 10 min. Protein samples were separated on Bolt Bis-Tris Plus Protein 4-12% gels (Thermo Scientific; #NW04120BOX) and then transferred to a nitrocellulose membrane using iBlot Transfer Stacks (Thermo Scientific; #IB23001). The membranes were then blocked in 5% milk or 5% BSA (for phospho-proteins) dissolved in TBST, incubated in primary antibodies overnight at 4°C, and secondary antibodies were applied the next day for 1 hr at room temperature. Bound antibodies were visualized using enhanced chemiluminescence detection reagents [Thermo Scientific; #34076 (West Dura), #A38554 (West Atto)] and imaged using the iBright CL750 Imaging System (Thermo Scientific; #A44116). Densitometric analysis was performed using NIH ImageJ software. Primary antibodies used for immunoblotting were goat-anti-VCAM-1 (1:1000, R&D Systems; #AF643), goat-anti-ICAM-1 (1:2000, R&D Systems; #AF796), rabbit-anti-NF-κB p65 (1:6000, Cell Signaling; #8242), rabbit-anti-phospho-NF-κB p65 (1:2000, Cell Signaling; #3033) and rabbit-anti-β-actin (1:4000, Cell Signaling; #4967). Secondary antibodies used were anti-rabbit-HRP (1:4000, Sigma; #A6667) and anti-goat-HRP (1:4000, Sigma; #A5420).

### Real-time quantitative reverse transcription PCR

Total RNA was extracted using TRIzol Reagent (Thermo Scientific; #15596018) and RNeasy Mini Kit (Qiagen; #74106). cDNA was prepared using an iScript cDNA Synthesis Kit (Bio-Rad; #1708891), and quantitative reverse transcription PCR was performed using SsoAdvanced Universal SYBR Green Supermix (Bio-rad; #1725274), gene primers (see **Supplemental Table S2**), and a CFX96 Touch Real-Time PCR System (Bio-Rad) or CFX Opus 384 Real-Time PCR System (Bio-Rad). Target genes were normalized against at least two of the following housekeeping genes: *Actb, Rn18s, Rpl13a, and Eife3*.

### BioRender

BioRender.com was used for generating the graphical abstract and Fig. 1C.

### Statistical analysis

All statistical tests were performed using GraphPad Prism software 9.0. Results are expressed as mean values ± standard error of the mean (SEM) unless indicated otherwise. All measurements were taken from distinct samples, and the same sample was not measured repeatedly. Comparisons between two groups were performed with a two-tailed Student’s *t*-test or with a one sample *t* and Wilcoxon test when data did not follow a normal distribution. Comparisons between more than two groups were made by one-way ANOVA followed by post-hoc comparisons. A *P* value of less than or equal to 0.05 was considered significant. Outlier tests were performed in Prism, but no data were excluded in the final graphs.

## Data availability

All data supporting the findings of this study are available within the manuscript and its Supplemental Materials. A list of all primers used for qRT-PCR and genotyping is provided in Supplemental Table S2. The Major Resources Table that is provided within the Supplemental Materials lists all mouse strains, primary and secondary antibodies, and the cell line used in this study.

## AUTHOR CONTRIBUTIONS

CFJ and CTG designed the study with significant scientific input from KYB, HC, and BGC. CFJ, CMS and KYB conducted experiments and acquired data. CFJ, CMS KYB, and BGC analyzed data. CFJ and CTG wrote the manuscript, which was critically revised and then approved by all authors before submission.

## ACKNOWLEDGEMENTS

We thank Dr. Jun Xie for technical assistance and members of the Griffin laboratory for valuable advice and feedback on this manuscript. We also thank Drs. Florea Lupu and Ravi Keshari for their guidance and technical advice in cytokine assays and Drs. Scott and Kendra Plafker for sharing the NRF2 antibody and for their assistance in NRF2 imaging and quantification. The graphical abstract and Figure 1C were created using BioRender.com.

