## Supplemental Materials for "Endothelial RIPK3 minimizes organotypic inflammation and vascular permeability in ischemia-reperfusion injury"

**SUPPLEMENTAL MATERIAL**
Supplemental Figures S1-S7
Supplemental Tables S1-2
Major Resources Table
ARRIVE Guidelines Table
Supplemental References

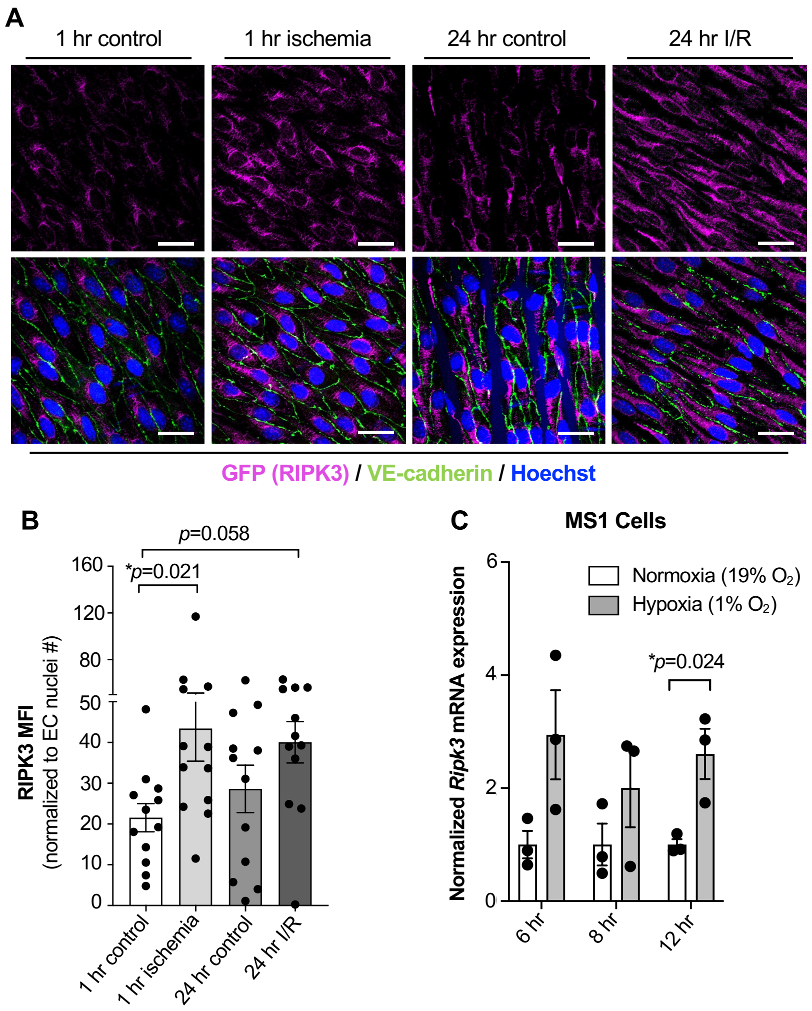

**SUPPLEMENTAL FIGURE S1:** ***Ischemia-reperfusion (I/R) injury and oxygen glucose deprivation increases RIPK3 expression in endothelial cells (ECs).*** (**A**) Mesenteric arterial EC RIPK3 protein expression was assessed by immunolabeling for GFP in RIPK3-GFP mice after 1 hr ischemia and after 24 hr intestinal I/R injury. 1 hr control and 24 hr control groups represent mesenteric arterial vessels from unclamped adjacent vessels. Scale bar: 20 µm. (**B**) Quantification of RIPK3 (GFP) expression after 1 hr ischemia and 24 hr I/R injury (n=4 mice/group, 3 representative images from each mouse). (**C**) *Ripk3* transcript expression in MS1 cells grown in glucose-free media and exposed to normoxia or hypoxia for the indicated time points (n=3/group). *P* values were determined by one-way ANOVA with posthoc comparisons (**B**) or unpaired t tests (**C**) for each treatment condition between groups. *$P < 0.05$ *vs*. Control. Summary data are the mean ± SE**. MFI** = mean fluorescence intensity

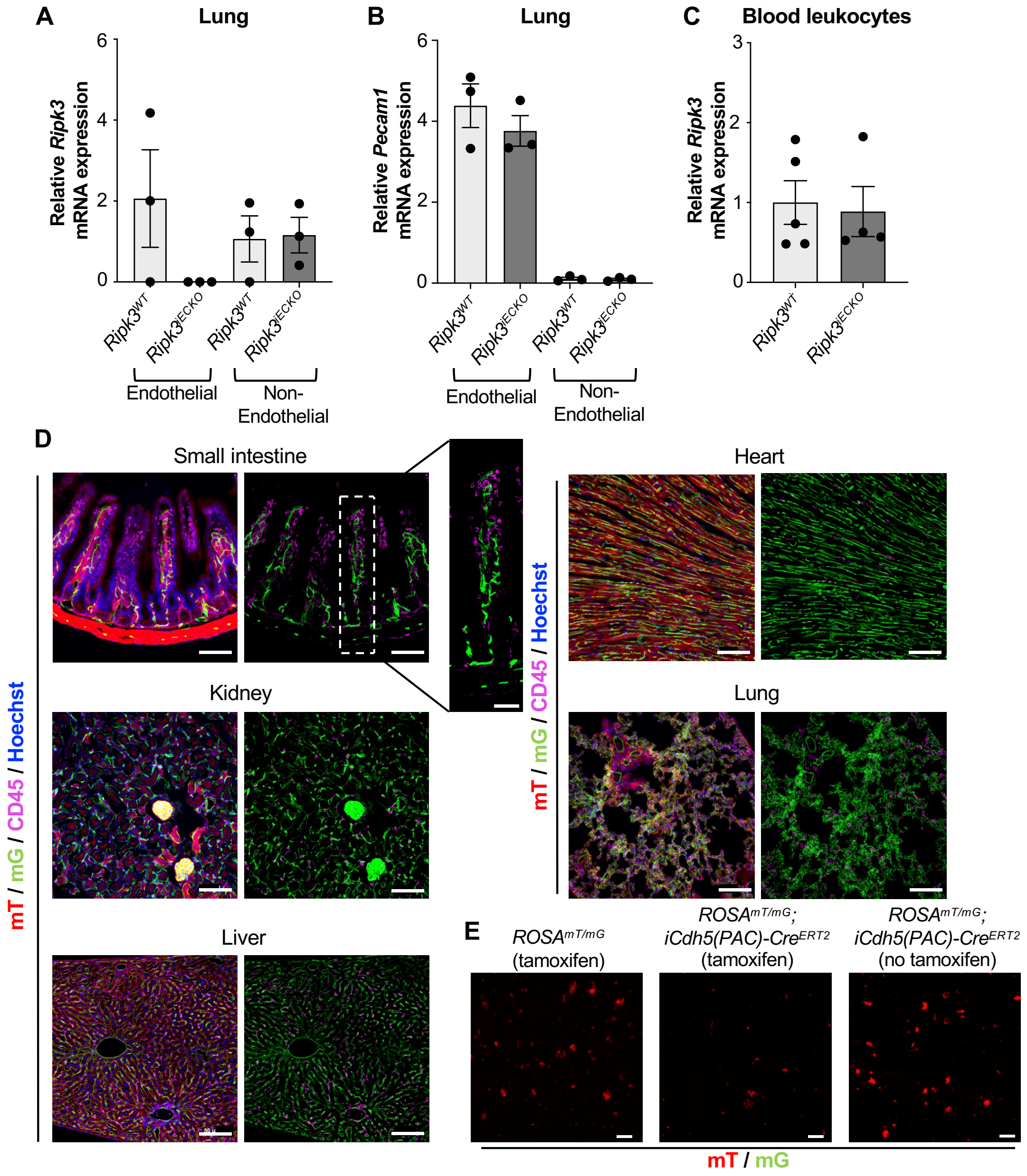

**SUPPLEMENTAL FIGURE S2: *Ripk3-flox deletion using the inducible Cdh5(PAC)-Cre$^{ERT2}$ transgenic line is specific to endothelial cells (ECs) in the adult mouse.*** (**A**) *Ripk3* transcript expression was assessed in ECs versus non-ECs enriched from lungs of *Ripk3*$^{WT}$ and *Ripk3*$^{iECKO}$ mice (n=3 mice/group). (**B**) *Pecam1* transcript expression in the same cellular compartments shown in (A) was assessed to indicate EC versus non-EC separation efficiency in lungs from *Ripk3*$^{WT}$ and *Ripk3*$^{iECKO}$ mice. (**C**) *Ripk3* transcript expression in circulating peripheral blood leukocytes from *Ripk3*$^{WT}$ and *Ripk3*$^{iECKO}$ mice (n=4-5 mice/group). (**D**) Cre activity (in green) was assessed in the indicated organs using confocal imaging after labeling for CD45$^{+}$ leukocytes in *ROSA*$^{mT/mG}$*;iCdh5(PAC)-Cre*$^{ERT2}$ mice. Scale bar: 100 µm. Enlarged image scale bar: 50 µm. (**E**) Cre activity (in green) was assessed in peripheral blood using *ROSA*$^{mT/mG}$ (negative control) or

*ROSA*$^{mT/mG}$*;iCdh5(PAC)-Cre*$^{ERT2}$ mice. Scale bar: 20 μm. *P* values were determined by one-sample t and Wilcoxon test (**A, endothelial cells**) or unpaired t-tests (**A, non-endothelial cells**; **B** and **C**) for each treatment condition between *Ripk3*$^{WT}$ and *Ripk3*$^{iECKO}$ groups. No *P* values reached significance, as defined by *$P$ <0.05 *vs*. Control. Summary data are the mean ± SE.

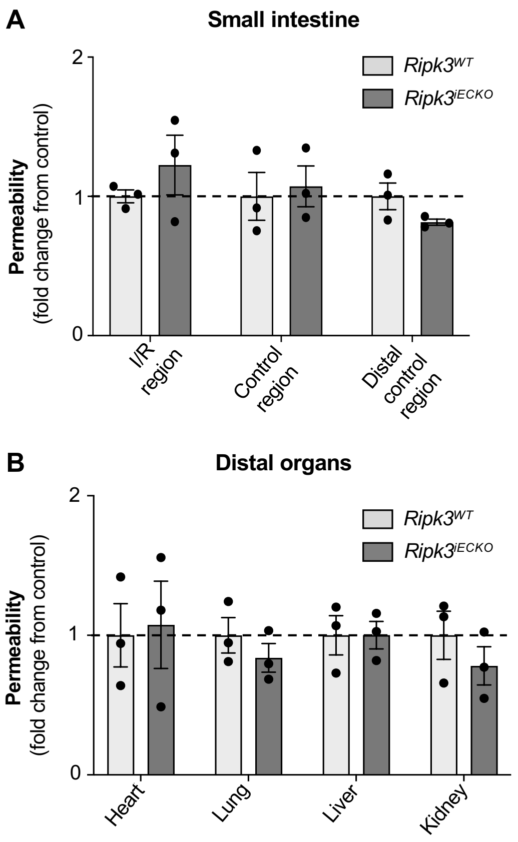

**SUPPLEMENTAL FIGURE S3:** ***Sham intestinal ischemia-reperfusion (I/R) injury operations do not cause permeability in Ripk3$^{WT}$ and Ripk3$^{iECko}$ mice.*** (**A**) Small intestinal vascular permeability measured by Evans blue dye leakage after 24 hr sham intestinal I/R injury in the injury region (I/R) and control regions (control, distal control) (n=3/group). (**B**) Vascular permeability in the indicated organs after 24 hr sham intestinal I/R injury (n=3/group). *P* values were determined by unpaired t-tests for each organ between the two groups **(A** and **B). ** No *P* values reached significance, as defined by $^*P < 0.05$ *vs*. Control. Summary data are the mean ± SE.

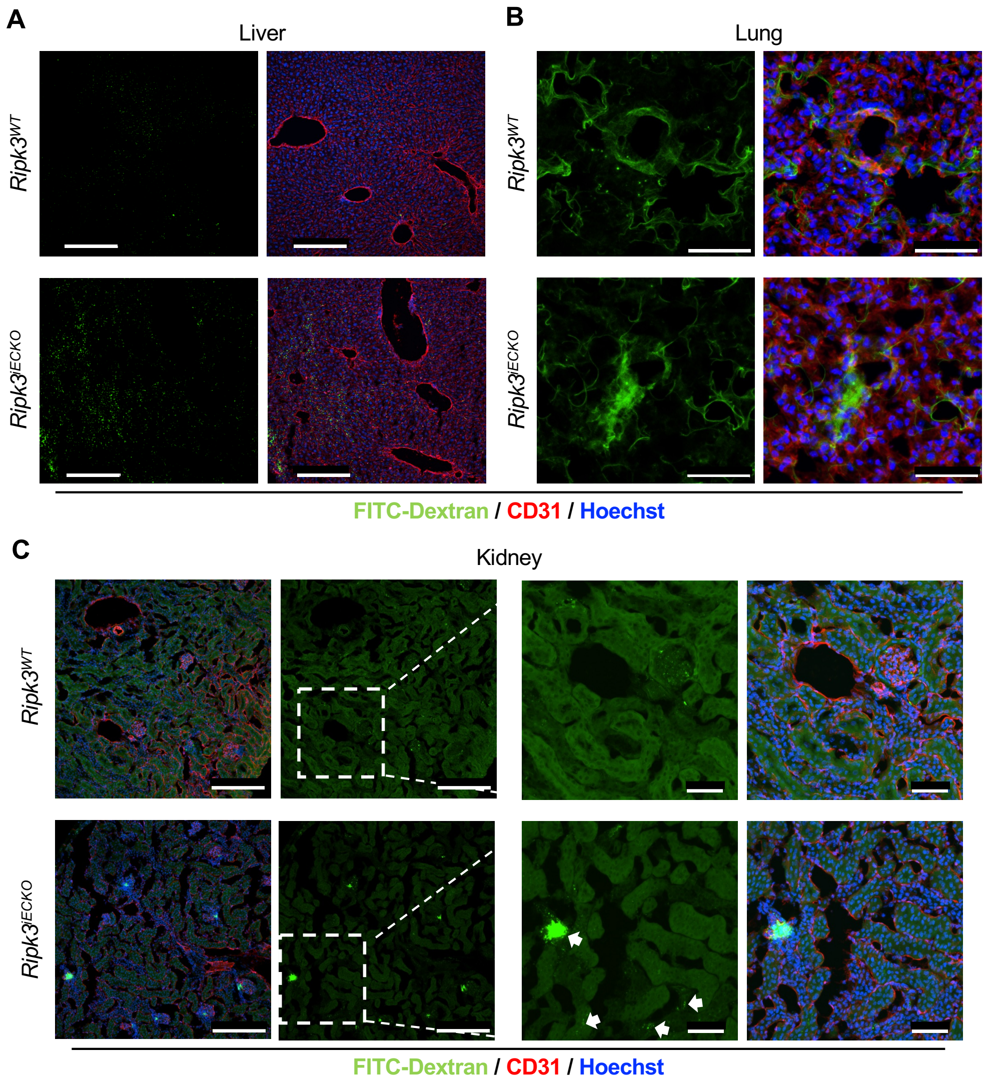

**SUPPLEMENTAL FIGURE S4:** ***FITC-Dextran vascular permeability following intestinal ischemia-reperfusion (I/R) injury in Ripk3$^{WT}$ and Ripk3$^{iECko}$ mice.*** Representative images of FITC-dextran (70kDa) permeability after 24 hr intestinal I/R injury in (**A**) livers, Scale bar: 200 µm, (**B**) lungs, Scale bar: 50 µm, and (**C**) kidneys with white arrows indicating permeability in glomeruli and peritubular capillaries in *Ripk3$^{iECKO}$* mice, Scale bar: 200 µm (left) and 50 µm for insets at right**.**

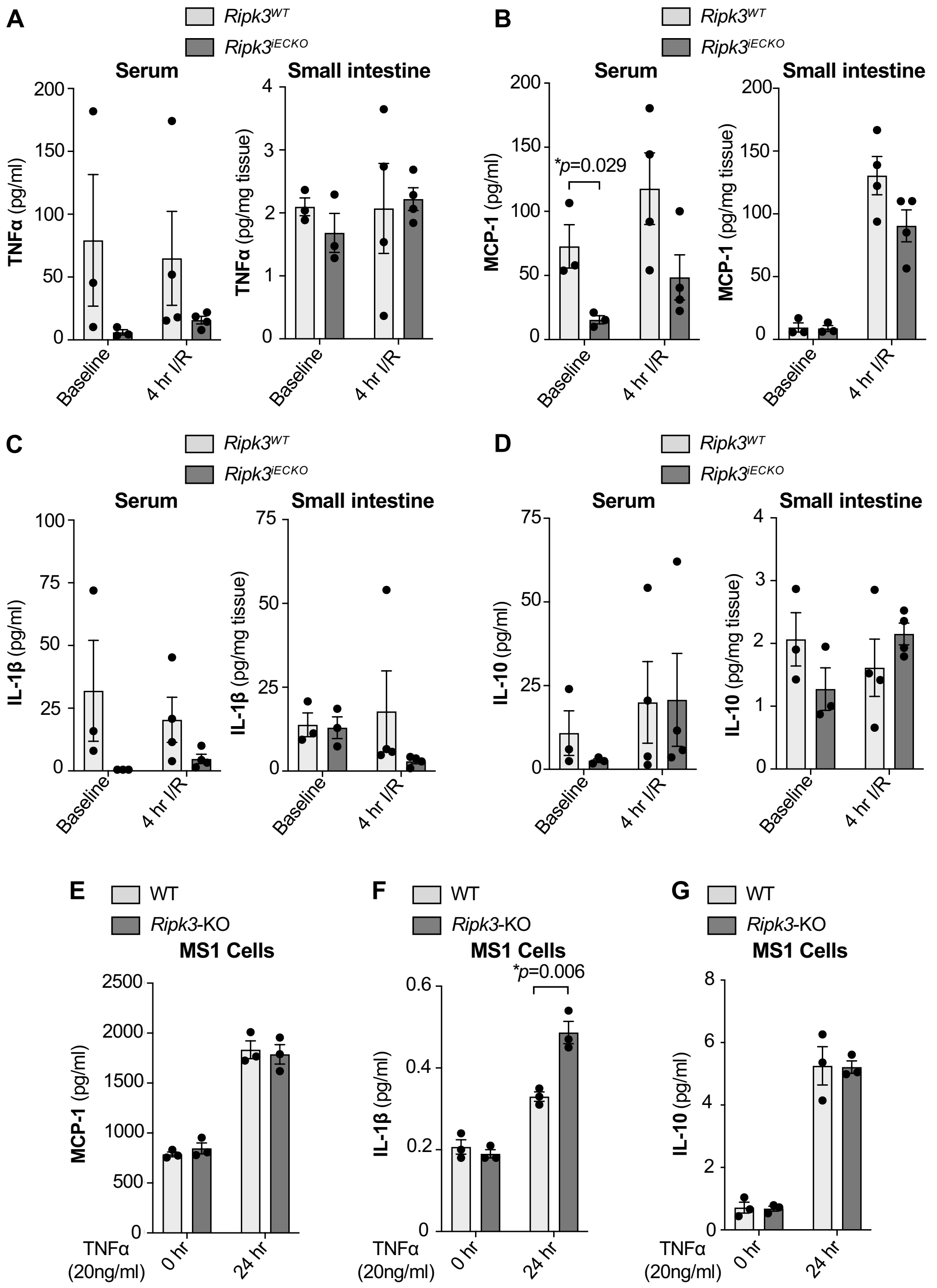

**SUPPLEMENTAL FIGURE S5:** ***Key cytokines not altered with endothelial Ripk3 deletion.*** Serum and small intestinal tissue (**A**) TNFα, (**B**) MCP-1, (**C**) IL-1β, and (**D**) IL-10 levels at baseline and following 4 hr ischemia/reperfusion (I/R) injury in *Ripk3*$^{WT}$ and *Ripk3*$^{iECKO}$ mice (n=3-4/group). (**E**) MCP-1, (**F**) IL-1β, and (**G**) IL-10 levels in MS1 endothelial cell supernatants after stimulating WT and *Ripk3*-KO cells with 20ng/ml TNFα for 0 hr or 24 hr (n=3/group)***.*** *P* values were determined by unpaired t-tests for each treatment condition between the two groups (**A-G**), and data that failed an equal variance F-test were log-transformed before analysis (**A, C,** and **D**). **P* <0.05 *vs*. Control. Summary data are the mean ± SE.

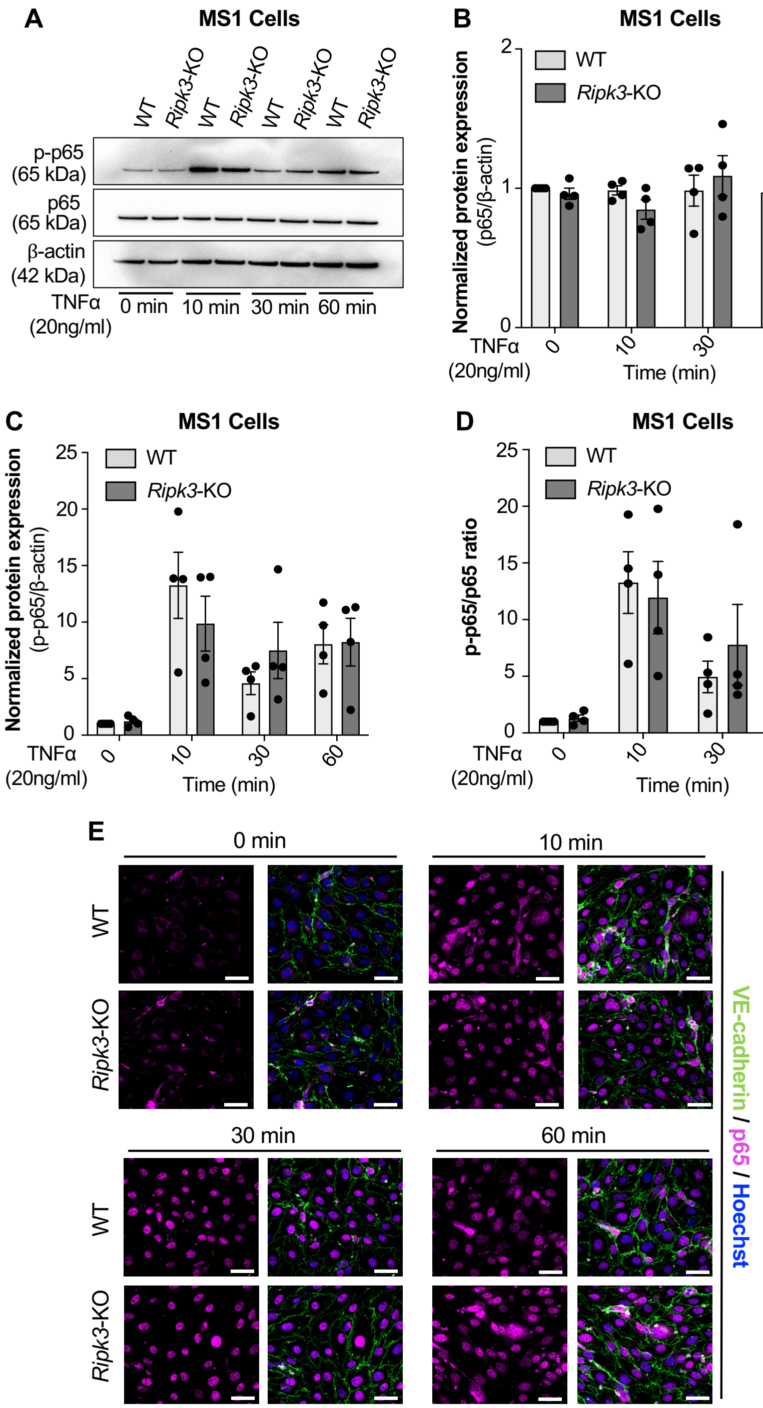

**SUPPLEMENTAL FIGURE S6:** ***NF-κB signaling and localization do not appear to be dysregulated in MS1 endothelial cells (ECs) with Ripk3 deletion.*** (**A**) Protein lysates from WT and *Ripk3*-KO MS1 ECs were immunoblotted for phospho-p65 (p-p65) and p65 after stimulation with 20ng/mL TNFα for the indicated times. (**B**) Quantified p65 protein expression between WT and *Ripk3*-KO MS1 ECs after TNFα treatment, as in A. (**C**) Quantified p-p65 protein expression between WT and *Ripk3*-KO cells after TNFα treatment, as in A. (**D**) p-p65/p65 ratio from data shown in B and C (n=4/group for A-D)**.** (**E**) WT and *Ripk3*-KO MS1 ECs were immunolabeled for p65 after stimulating with 20ng/mL TNFα for the indicated times, Scale bar: 20 μm. *P* values were determined by unpaired t-tests for each treatment timepoint between the two groups (**B**, **C** and **D**). No *P* values reached significance, as defined by *$P$ <0.05 *vs*. Control. Summary data are the mean ± SE.

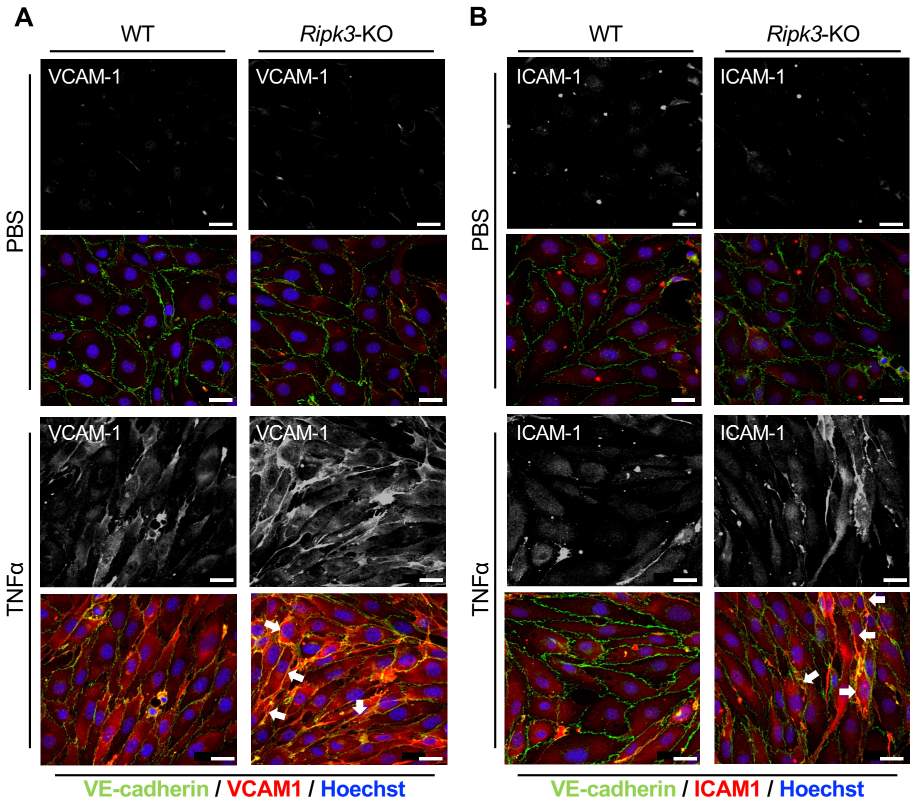

**SUPPLEMENTAL FIGURE S7:** ***Increased vascular cell adhesion molecule-1 (VCAM-1) and intercellular adhesion molecule-1 (ICAM-1) expression in Ripk3-KO MS1 endothelial cells (ECs).*** WT and *Ripk3*-KO MS1 ECs were stimulated with either PBS or 20ng/mL TNFα for 24 hr and immunostained for VCAM-1 (**A**) or ICAM-1 (**B**). White arrows indicate junctional localization of VCAM-1 or ICAM-1, Scale bar: 20 µm.

**SUPPLEMENTAL TABLE S1:** ***Mouse sexes used in individual experiments***

| Figure | Condition | Males | Females |
|---|---|---|---|
| **1B** | *Ripk3*$^{WT}$ mice: organ baseline permeability | 3 | 2 |
| | *Ripk3*$^{iECKO}$ mice: organ baseline permeability | 3 | 2 |
| **1C and 1D** | *Ripk3*$^{WT}$ mice: organ permeability after 24 hr I/R injury | 4 | 2 |
| | *Ripk3*$^{iECKO}$ mice: organ permeability after 24 hr I/R injury | 4 | 2 |
| **2A and 2B and S5A-D** | *Ripk3*$^{WT}$ mice: baseline serum/small intestinal IL-6 levels | 1 | 2 |
| | *Ripk3*$^{iECKO}$ mice: baseline serum/small intestinal IL-6 levels | 1 | 2 |
| | *Ripk3*$^{WT}$ mice: serum/small intestinal IL-6 levels after 4 hr I/R injury | 3 | 1 |
| | *Ripk3*$^{iECKO}$ mice: serum/small intestinal IL-6 levels after 4 hr I/R injury | 1 | 3 |
| **2C and 2D** | Control mice: plasma IL-6 levels at 2 hr or 4 hr after PBS and LPS stimulation in whole blood | 2 | 1 |
| | *Ripk3*$^{iECKO}$ mice: plasma IL-6 levels at 2 hr or 4 hr after PBS and LPS stimulation in whole blood | 3 | 0 |
| **4A** | *Ripk3*$^{WT}$ mice: baseline small intestinal VCAM-1 expression | 3 | 4 |
| | *Ripk3*$^{iECKO}$ mice: baseline small intestinal VCAM-1 expression | 2 | 5 |
| **4B** | *Ripk3*$^{WT}$ mice: baseline lung VCAM-1 expression | 4 | 2 |
| | *Ripk3*$^{iECKO}$ mice: baseline lung VCAM-1 expression | 4 | 2 |
| **4B** | *Ripk3*$^{WT}$ mice: baseline liver VCAM-1 expression | 2 | 4 |
| | *Ripk3*$^{iECKO}$ mice: baseline liver VCAM-1 expression | 2 | 4 |
| **4B** | *Ripk3*$^{WT}$ mice: baseline kidney VCAM-1 expression | 4 | 2 |
| | *Ripk3*$^{iECKO}$ mice: baseline kidney VCAM-1 expression | 4 | 2 |
| **5A** | *Ripk3*$^{WT}$ mice: baseline lung ICAM-1 expression | 4 | 2 |

| | | | |
|---|---|---|---|
| | *Ripk3*$^{iECKO}$ mice: baseline lung ICAM-1 expression | 4 | 2 |
| **5B** | *Ripk3*$^{WT}$ mice: baseline small intestinal ICAM-1 expression | 3 | 4 |
| | *Ripk3*$^{iECKO}$ mice: baseline small intestinal ICAM-1 expression | 2 | 5 |
| **5B** | *Ripk3*$^{WT}$ mice: baseline liver ICAM-1 expression | 2 | 4 |
| | *Ripk3*$^{iECKO}$ mice: baseline liver ICAM-1 expression | 2 | 4 |
| **5B** | *Ripk3*$^{WT}$ mice: baseline kidney ICAM-1 expression | 4 | 2 |
| | *Ripk3*$^{iECKO}$ mice: baseline kidney ICAM-1 expression | 4 | 2 |
| **7B** | Control mice: Peripheral blood monocyte numbers after PBS liposome administration | 3 | 2 |
| | Control mice: Peripheral blood monocyte numbers after clodronate liposome administration | 4 | 2 |
| **7D and 7E** | *Ripk3*$^{WT}$ mice: organ 24 hr I/R permeability after clodronate induced monocyte depletion | 2 | 3 |
| | *Ripk3*$^{iECKO}$ mice: organ 24 hr I/R permeability after clodronate induced monocyte depletion | 2 | 2 |
| **S1B** | *Ripk3-GFP* mice: RIPK3 expression after 1 hr ischemia | 1 | 3 |
| | *Ripk3-GFP* mice: RIPK3 expression after 24 hr I/R injury | 1 | 3 |
| **S2A and S2B** | *Ripk3*$^{WT}$ mice: lung *Ripk3* and *Pecam1* mRNA expression | 2 | 1 |
| | *Ripk3*$^{iECKO}$ mice: lung *Ripk3* and *Pecam1* mRNA expression | 2 | 1 |
| **S2C** | *Ripk3*$^{WT}$ mice: peripheral blood leukocyte *Ripk3* and *Pecam1* mRNA expression | 5 | 0 |
| | *Ripk3*$^{iECKO}$ mice: peripheral blood leukocyte *Ripk3* and *Pecam1* mRNA expression | 4 | 0 |
| **S3A and S3B** | *Ripk3*$^{WT}$ mice: organ permeability after sham I/R injury | 2 | 1 |
| | *Ripk3*$^{iECKO}$ mice: organ permeability after sham I/R injury | 1 | 2 |

**SUPPLEMENTAL TABLE S2:** ***List of all primers used for quantitative reverse transcription PCR (qRT-PCR) and genotyping***

| Primer Name | Sequence | Application |
|---|---|---|
| ***Ripk3*-F** | 5'-CCTTCCAGGACTGCGAACCA | qRT-PCR: *Ripk3* expression |
| ***Ripk3*-R** | 5'-TGCCTCTTTGGCTTGGCTCT | |
| ***Vcam1*-F** | 5'-TGAACCCAAACAGAGGCAGAGT | qRT-PCR: *Vcam1* expression |
| ***Vcam1*-R** | 5'-GGTATCCCATCACTTGAGCAGG | |
| ***Icam1*-F** | 5'-CAATTTCTCATGCCGCACAG | qRT-PCR: *Icam1* expression |
| ***Icam1*-R** | 5'-AGCTGGAAGATCGAAAGTCCG | |
| ***Il6*-F** | 5'-CAAAGCCAGAGTCCTTCAGAG | qRT-PCR: *Il6* expression |
| ***Il6*-R** | 5'-TGGTCCTTAGCCACTCCTTC | |
| ***Pecam1-F*** | 5'-AGTCAGAGTCTTCCTTGCCC | qRT-PCR: *Pecam1* expression |
| ***Pecam1-R*** | 5'-TCTGTTTGGCCTTGGCTTTC | |
| ***Rpl13a*-F** | 5'-GAGGTCCTCAAGACCAACGG | qRT-PCR: *Ripl13a* expression |
| ***Rpl13a*-R** | 5'-GGTACTTGGTCAGGTGGGTG | |
| ***Eif3e*-F** | 5'-GGTTGGATGCCAAGATTGATTC | qRT-PCR: *Eif3e* expression |
| ***Eif3e*-R** | 5'-GGGCGAGACTGCATTGTTG | |
| ***Actb*-F** | 5'-TGTTACCAACTGGGACGACA | qRT-PCR: *Actb* expression |
| ***Actb*-R** | 5'-GGGGTGTTGAAGGTCTCAAA | |
| ***Rn18s*-F** | 5'-CCCGAAGCGTTTACTTTGAAA | qRT-PCR: *Rn18s* expression |
| ***Rn18s*-R** | 5'-CGCGGTCCTATTCCATTATTC | |
| ***Nfe2l2*-F** | 5'-ACTACAGTCCCAGCAGGACAT | qRT-PCR: *Nfe2l2* expression |
| ***Nfe2l2*-R** | 5'-CTGTTCCTTCTGGAGTTGCTCTT | |
| ***Hmox1*-F** | 5'-GACAGCCCCACCAAGTTCAA | |

| | | |
|---|---|---|
| ***Hmox1*-R** | 5’-AGCTCCTCAAACAGCTCAATGT | qRT-PCR: *Hmox1* expression |
| ***Nqo1*-F** | 5’-GGCCGATTCAGAGTGGCAT | qRT-PCR: *Nqo1* expression |
| ***Nqo1*-R** | 5’-CCAGACGGTTTCCAGACGTT | |
| ***Ripk3-flox*-F** | 5’-CCATCCTCCCTTCATCAAAA | Genotyping: *Ripk3-flox* |
| ***Ripk3-flox*-R** | 5’-CGGACTTTGAATGAGCGACT | |
| ***Ripk3-wt*-F** | 5’-ACCTCTGAACCCTCTCACCA | Genotyping: *Ripk3-gfp* |
| ***Ripk3-gfp*-F** | 5’-AGCGCGATCACATGGTCCTG | |
| ***Ripk3*-R** | 5’-TCTTAGCCAGTCCAGGTTGC | |
| ***iCdh5-Cre*-F** | 5’-TCCTGATGGTGCCTATCCTC | Genotyping: *iCdh5-Cre*$^{ERT2}$ |
| ***iCdh5-Cre*-R** | 5’-CGAACCTGGTCGAAATCAGT | |
| **Control-F** | 5’-CTAGGCCACAGAATTGAAAGATCT | |
| **Control-R** | 5’-GTAGGTGGAAATTCTAGCATCATCC | |
| ***mTmG*-F** | 5’-CTCTGCTGCCTCCTGGCTTCT | Genotyping: *Rosa*$^{mTmG}$ |
| ***mTmG*-R** | 5’-CGAGGCGGATCACAAGCAATA | |
| ***mTmG*-R** | 5’-TCAATGGGCGGGGGTCGTT | |

**MAJOR RESOURCES TABLE**

### Animals (in vivo studies)

| Species | Vendor or Source | Background Strain | Sex | Persistent ID / URL |
|---|---|---|---|---|
| Mouse | Our Laboratory | *Ripk3$^{fl/fl}$* | M/F | Original Publication: [1] |
| Mouse | Taconic | *Cdh5(PAC)-Cre$^{ERT2}$* | M/F | #13073 |
| Mouse | Jackson Laboratory | *Ripk3-gfp$^{fl/fl}$* | M/F | RRID:IMSR_JAX:030284 |
| Mouse | Jackson Laboratory | *ROSA$^{mT/mG}$* | M/F | RRID:IMSR_JAX:007576 |

### Primary Antibodies

| Target antigen | Vendor or Source | Catalog # | Working concentration | Persistent ID / URL |
|---|---|---|---|---|
| GFP | Novus | NB600-303 | 1:200 | RRID:AB_523899 |
| VE-cadherin | Abcam | ab91064 | 1:100 | RRID:AB_2049374 |
| CD45 | R&D Systems | AF114 | 1:50 | RRID:AB_442146 |
| F4/80 | Cell Signaling | 30325 **(R)** | 1:300 | RRID:AB_2798990 |
| VCAM-1 | R&D Systems | AF643 | 1:100 – 1:1000 | RRID:AB_355499 |
| ICAM-1 | R&D Systems | AF796 | 1:100 – 1:2000 | RRID:AB_2248703 |
| p65 (NF-κB) | Cell Signaling | 8242 **(R)** | 1:700 – 1:6000 | RRID:AB_10859369 |
| Phospho-p65 (NF-κB) | Cell Signaling | 3033 **(R)** | 1:2000 | RRID:AB_331284 |
| β-actin | Cell Signaling | 4967 | 1:4000 | RRID:AB_330288 |
| CD31 | R&D Systems | AF3628 | 1:100 | RRID:AB_2161028 |
| CD31 | BD Pharmingen | 553370 | 1:100 | RRID:AB_394816 |
| NRF2 | Gift from Plafker Laboratory at OMRF | | 1:300 | Original Publication: [2] |

### Secondary Antibodies

| Host and Target species | Fluorophore or conjugate | Vendor or Source | Catalog # | Working concentration | Persistent ID / URL |
|---|---|---|---|---|---|

| | | | | | |
|---|---|---|---|---|---|
| Donkey anti-Rabbit | Cy5 | Jackson ImmunoResearch | 711-175-152 | 1:500 | RRID:AB_2340607 |
| Donkey anti-Rat | Alexa488 | Thermo Fisher | A-21208 | 1:500 | RRID:AB_2535794 |
| Donkey anti-Goat | Cy3 | Jackson ImmunoResearch | 705-165-003 | 1:500 | RRID:AB_2340411 |
| Goat anti-Rabbit | HRP | Sigma | A6667 | 1:4000 | RRID:AB_258307 |
| Rabbit anti-Goat | HRP | Sigma | A5420 | 1:4000 | RRID:AB_258242 |

**Cultured Cells**

| Name | Vendor or Source | Sex (F, M, or unknown) | Persistent ID / URL |
|---|---|---|---|
| MS1 pancreatic islet endothelial cells | ATCC | Unknown | CRL-2279 |

**ARRIVE GUIDELINES TABLE**

**Figure 1B**

| Groups | Sex | Age | Number (prior to experiment) | Number (after termination) | Littermates (Yes/No) | Other description |
|---|---|---|---|---|---|---|
| *Ripk3*$^{WT}$ | M/F | 12 weeks old | 5 | 5 | Littermates from two litters | |
| *Ripk3*$^{iECKO}$ | M/F | 12 weeks old | 5 | 5 | Littermates from two litters | |

**Figure 1C-D**

| Groups | Sex | Age | Number (prior to experiment) | Number (after termination) | Littermates (Yes/No) | Other description |
|---|---|---|---|---|---|---|
| *Ripk3*$^{WT}$ | M/F | 12 weeks old | 6 | 6 | Littermates from two litters | |
| *Ripk3*$^{iECKO}$ | M/F | 12 weeks old | 6 | 6 | Littermates from two litters | |

**Figure 2A-B and Figure S5A-D**

| Groups | Sex | Age | Number (prior to experiment) | Number (after termination) | Littermates (Yes/No) | Other description |
|---|---|---|---|---|---|---|
| *Ripk3*$^{WT}$ (baseline) | M/F | 12 weeks old | 3 | 3 | Yes | |
| *Ripk3*$^{iECKO}$ (baseline) | M/F | 12 weeks old | 3 | 3 | Yes | |

| | | | | | |
|---|---|---|---|---|---|
| *Ripk3*$^{WT}$ (4 hr I/R injury) | M/F | 12 weeks old | 4 | 4 | Yes |
| *Ripk3*$^{iECKO}$ (4 hr I/R injury) | M/F | 12 weeks old | 4 | 4 | Yes |

**Figure 2C-D**

| Groups | Sex | Age | Number (prior to experiment) | Number (after termination) | Littermates (Yes/No) | Other description |
|---|---|---|---|---|---|---|
| *Ripk3*$^{WT}$ | M/F | 12 weeks old | 3 | 3 | Yes | |
| *Ripk3*$^{iECKO}$ | M | 12 weeks old | 3 | 3 | Yes | |

**Figure 4A-B and 5A-B**

| Groups | Sex | Age | Number (prior to experiment) | Number (after termination) | Littermates (Yes/No) | Other description |
|---|---|---|---|---|---|---|
| *Ripk3*$^{WT}$ | M/F | 12 weeks old | 6-7 | 6-7 | Littermates from two litters | |
| *Ripk3*$^{iECKO}$ | M/F | 12 weeks old | 6-7 | 6-7 | Littermates from two litters | |

**Figure 7B**

| Groups | Sex | Age | Number (prior to experiment) | Number (after termination) | Littermates (Yes/No) | Other description |
|---|---|---|---|---|---|---|
| Control-PBS liposomes | M/F | 12 weeks old | 5 | 5 | Littermates from two litters | |

| | | | | | |
|---|---|---|---|---|---|
| Control-Clodronate liposomes | M/F | 12 weeks old | 6 | 6 | Littermates from two litters |

**Figure 7D-E**

| Groups | Sex | Age | Number (prior to experiment) | Number (after termination) | Littermates (Yes/No) | Other description |
|---|---|---|---|---|---|---|
| *Ripk3*$^{WT}$ | M/F | 12 weeks old | 6 | 5 | Littermates from two litters | |
| *Ripk3*$^{iECKO}$ | M/F | 12 weeks old | 5 | 4 | Littermates from two litters | |

**Figure S1B**

| Groups | Sex | Age | Number (prior to experiment) | Number (after termination) | Littermates (Yes/No) | Other description |
|---|---|---|---|---|---|---|
| RIPK3-GFP 1 hr ischemia only | M/F | 10-12 weeks old | 4 | 4 | Yes | |
| RIPK3-GFP 24 hr I/R injury | M/F | 10-12 weeks old | 4 | 4 | Yes | |

**Figure S2A-B**

| Groups | Sex | Age | Number (prior to experiment) | Number (after termination) | Littermates (Yes/No) | Other description |
|---|---|---|---|---|---|---|
| *Ripk3*$^{WT}$ | M/F | 12 weeks old | 3 | 3 | Yes | |

| | | | | | |
|---|---|---|---|---|---|
| *Ripk3$^{iECKO}$* | M/F | 12 weeks old | 3 | 3 | Yes |

**Figure S2C**

| Groups | Sex | Age | Number (prior to experiment) | Number (after termination) | Littermates (Yes/No) | Other description |
|---|---|---|---|---|---|---|
| *Ripk3$^{WT}$* | M | 12 weeks old | 5 | 5 | Littermates from two litters | |
| *Ripk3$^{iECKO}$* | M | 12 weeks old | 4 | 4 | Littermates from two litters | |

**Figure S3A-B**

| Groups | Sex | Age | Number (prior to experiment) | Number (after termination) | Littermates (Yes/No) | Other description |
|---|---|---|---|---|---|---|
| *Ripk3$^{WT}$* | M | 12 weeks old | 3 | 3 | Yes | |
| *Ripk3$^{iECKO}$* | M | 12 weeks old | 3 | 3 | Yes | |
